# A planar-polarized MYO6-DOCK7-RAC1 axis promotes tissue fluidification in mammary epithelia

**DOI:** 10.1101/2023.01.23.524898

**Authors:** Luca Menin, Janine Weber, Stefano Villa, Emanuele Martini, Elena Maspero, Valeria Cancila, Paolo Maiuri, Andrea Palamidessi, Emanuela Frittoli, Fabrizio Bianchi, Claudio Tripodo, Kylie J. Walters, Fabio Giavazzi, Giorgio Scita, Simona Polo

## Abstract

Tissue fluidification and collective motility are pivotal in regulating embryonic morphogenesis, wound healing and tumor metastasis. These processes frequently require that each cell constituent of a tissue coordinates its migration activity and directed motion through the oriented extension of lamellipodia cell protrusions, promoted by RAC1 activity. While the upstream RAC1 regulators in individual migratory cells or leader cells during invasion or wound healing are well characterized, how RAC1 is controlled in follower cells remains unknown. Here, we identify a novel MYO6-DOCK7 axis that is critical for spatially restriction of RAC1 activity in a planar polarized fashion in model tissue monolayers. The MYO6-DOCK7 axis specifically controls the extension of cryptic lamellipodia required to drive tissue fluidification and cooperative mode motion in otherwise solid and static carcinoma cell collectives.

**Highlights:**

- Collective motion of jammed epithelia requires myosin VI activity
- The MYO6-DOCK7 axis is critical to restrict the activity of RAC1 in a planar polarized fashion
- MYO6-DOCK7-RAC1 activation ensures long-range coordination of movements by promoting orientation and persistence of cryptic lamellipodia
- Myosin VI overexpression is exploited by infiltrating breast cancer cells

## INTRODUCTION

Collective motility and tissue fluidification are emerging as key regulators in physiological tissue remodeling during development, wound repair, and regeneration, and in pathological conditions, first and foremost during carcinoma dissemination [1-5]. During these processes, individual cells composing a given tissue coordinate their motion with that of their neighbor cells by keeping tight cell-cell contacts to migrate or invade [6].

A remarkable form of collective dynamics occurs during the early development of epithelial and glandular tissues, which are characterized by individual cells that constantly rearrange their motion, as in a fluid. This fluid-like property endows tissues with a large degree of plasticity that is instrumental in the initial phases of tissue specification and morphogenesis. As density rises due to proliferation and tissues mature and differentiate, the motion of each cell is constrained by the crowding of its neighbors, forcing cells to move in groups, in a highly coordinated and cooperative fashion [7, 8]. At a critical density, motility ceases and tissues rigidify to undergo a fluid-to-solid phase transition, a process recently referred to as a jamming transition [9]. This tissue-level phase transition has been proposed to be critical for the development of the elastic properties and barrier function of epithelial tissues and might also act as an intrinsic homeostatic mechanical barrier to the development and expansion of structurally altered, hyperdynamic oncogenic clones. Conversely, a certain degree of fluidity is needed for a tissue to repair its wound, proliferate, or locally disseminate, such as during carcinoma progression [1, 4].

Tissue fluidification and collective motility are ruled by biochemical and physical interactions that cells establish with each other and their environment [2, 10]. How cells and tissues regulate this process and control these parameters has only begun to be investigated. To drive and propel directed cell migration, cells need to dynamically reorganize their actin cytoskeleton and generate actively pushing lamellipodial cell protrusions [11]. When embedded within a fully confluent monolayer or a developing epithelial tissue, cell protrusions can no longer extend into free space, but are forced to either push adjacent neighboring cells or slip underneath them, and are commonly referred to as cryptic lamellipodia [12]. The coordinated and directed extension of cryptic lamellipodia along a common direction drives groups of cells to move either as a solid rotating flock, such as in the case of the follicular epithelial cells in *Drosophila melanogaster* [13], or as supracellular streams that fluidize the whole tissue, as during compressive-or endocytic-driven unjamming of human bronchial epithelial tissues and model mammary epithelial cells [14, 15].

We previously showed that elevated levels of RAB5A, a master regulator of early endosomes, is sufficient to re-awaken the motility of otherwise kinetically-arrested epithelial monolayers, promoting millimeter-scale flocking-fluid motility through large-scale coordinated migration and local cell rearrangements [14, 16-18]. At the molecular level, RAB5A impinges on junction topology, turnover and tension, fostering the extension of oriented and coordinated cryptic lamellipodia [14, 17]. The leading edges of cryptic lamellipodia, much like those of individual crawling cells, are generated by localized and spatially restricted actin polymerization triggered by numerous actin regulators and coordinated by the small GTPase RAC1 [14, 19]. However, the molecular machinery driving cryptic lamellipodia protrusion is likely to be distinct or enriched with specific components with respect to more canonical lamellipodia of cells moving individually or into free space. Indeed, a distinct cell identity with a leader-to-follower topological organization is emerging as critical in driving directed collective motion, such as in the case of Indian file moving cells [20], protruding multicellular fingers extending into the free space during wound closure in model epithelia [21], or cell cohort invading stromal tissues during carcinoma dissemination [4, 22, 23].

Among the proteins that regulate cell migration and membrane protrusion, myosin VI is an unconventional actin-based motor protein that moves toward the minus end of the actin filaments [24-26]. Myosin VI was originally characterized in *Drosophila melanogaster*, where it participates in the collective migration of border cells during ovary development [27, 28]. The pro-migratory function of myosin VI has been confirmed also in tumors where elevated levels of this protein are frequent and correlate with the aggressive behavior in ovarian, breast, and prostate cancer [26, 28-31]. In epithelial carcinoma, deregulated alternative splicing results in the preferential or exclusive expression of the short isoform of myosin VI, to which cancerous malignancy becomes addicted for migration [31]. Despite the increasing interest in the potential oncogenic role of myosin VI in cell migration, little is known about its mechanism of action and potential impact on cancer collective migratory capacity.

Here, we employ oncogenic monolayer models in which RAB5A expression can be tuned in an inducible fashion to promote flocking-fluid motility and test the impact of myosin VI. We uncover a myosin VI-DOCK7 axis as critical for spatially restricting the activity of RAC1 in a planar polarized fashion. Myosin VI is specifically required in the follower cells to promote the formation of cryptic lamellipodia that drive tissue fluidification by triggering highly coordinated and cooperative mode motion in otherwise solid and static carcinoma cell collectives.

## RESULTS

### Myosin VI is critical for the coherent motion of jammed epithelia

Myosin VI is required for cancer cell migration through an ill-defined molecular mechanism [25, 31]. We examined the role of myosin VI during both individual and collective cell migration; namely, during random and confined migration of single cells, or in wound healing and tissue fluidification via flocking stream in epithelial monolayers (Figure 1A). To directly compare the results, we exploited the oncogenically transformed MCF10.DCIS.com cell line in which collective streaming of cells in densely-packed and confluent monolayers can be induced by modulation of RAB5A expression ([14], hereafter referred to as DCIS-RAB5A). This cell line uniquely expresses the short isoform of myosin VI (Figure S1A) and, thus, represents the ideal model system to identify specific roles of this isoform.

**Figure 1.**
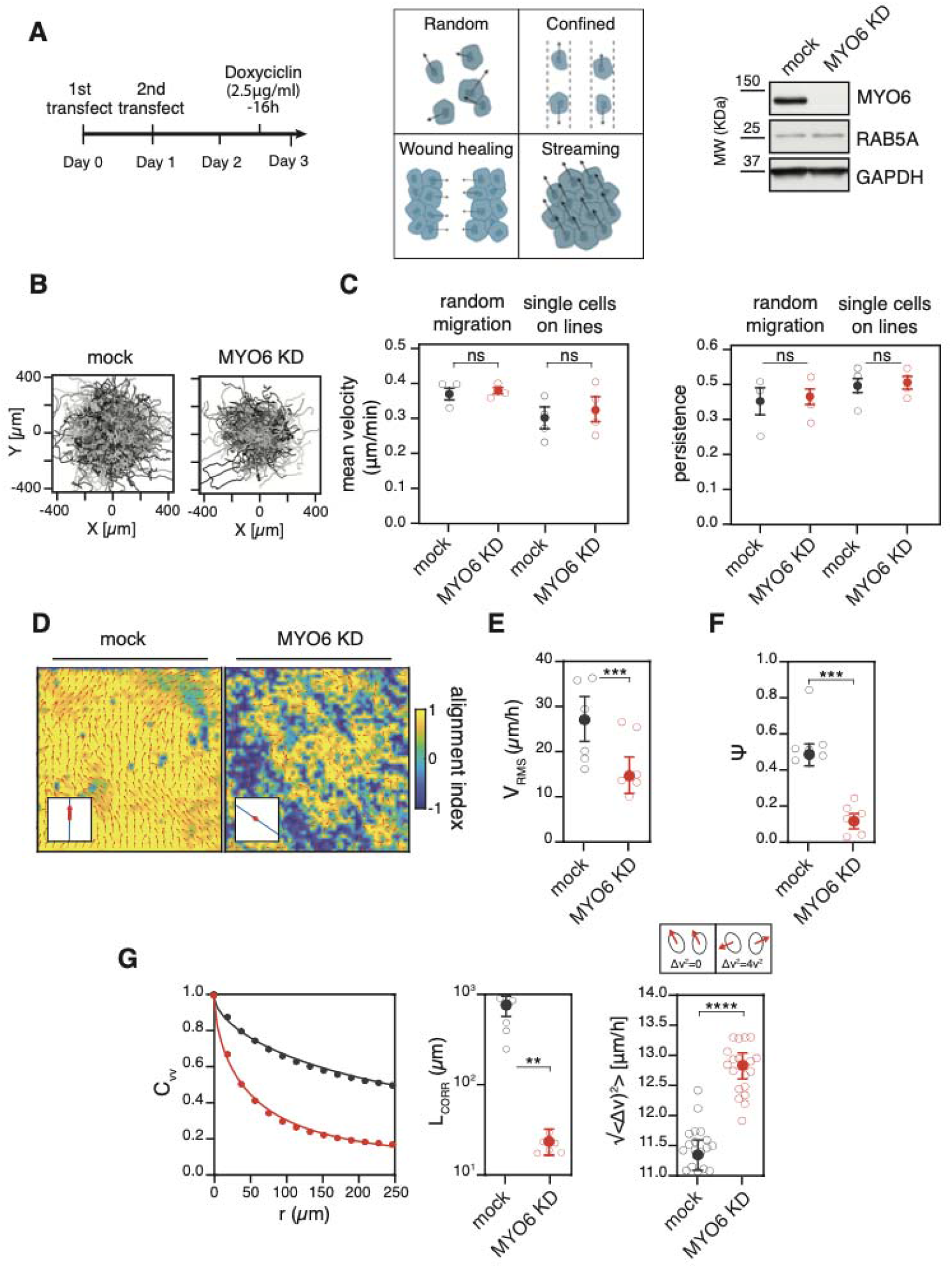
Myosin VI is required for coherent motion of jammed epithelia. **(A)** Experimental pipeline used for all kinetic experiments using DCIS-RAB5A and derivative cell lines. Cells were transfected twice with siRNA oligos, first in suspension and the following day in adhesion. 16 hours before the experiment, RAB5A expression was induced with 2.5L□g/mL doxycycline. Right panel, representative immunoblot (IB) analysis of the myosin VI depleted lysates performed at the end of the experiment (day 3 post siRNA oligos 1^st^ transfection). **(B)** Representative example of single cell trajectories obtained from random migration assays with the indicated cell lines. See also Video S1 and S2. **(C)** Quantification of single cell motion assays using DCIS-RAB5A H2B-mCherry mock and myosin VI depleted cell lines. Single cell mean velocity (left panel) and cell migration persistence (right panel) were quantified from random or confined migration assays, tracking the motion of tagged nuclei of the indicated genotype. *n* =75 (at least 25 single cells/experiment/genotype for at least three independent experiments). Empty circle, mean of the single experiment. Error bars, ±SD. ns > 0.999 by Student’s t-test. **(D)** Representative snapshots of the velocity field obtained by PIV analysis (red arrows), taken at 8 h after the beginning of the cell streaming experiment. DCIS-RAB5A H2B-mCherry mock or myosin VI depleted cells (KD) monolayers were monitored by time-lapse phase contrast microscopy for 24 hours (Video S4) and analyzed by PIV (Video S5) to determine the local prevalent direction of motion resulting in velocity field’s vectors. The color map represents the alignment with respect to the mean instantaneous velocity, quantified by the alignment index *a*(*x*) = ***v***(x) · ***v***_0_/|***v***(x) · ***v***_0_| that is equal to 1(−1) when the local velocity is parallel (antiparallel) to the mean direction of migration. Red arrows in each inset represent the mean velocity v_O_ (average over the entire field of view). **(E)** Root mean squared velocity *V*_*RMS*_ parameter obtained from the PIV analysis of six independent experiments. Empty circle, mean of each experiment calculated from at least five videos/condition. Error bars, ±SD. ***P < 0.001 by Student’s t-test. **(F)** Orientational order parameter *Ψ* obtained from the PIV analysis as in (E). Error bars, ±SD. ***P < 0.001 by Student’s t-test. **(G)** Velocity correlation function *C*_*vv*_(r) and the corresponding correlation length *L*_*corr*_ obtained from the PIV analysis. Error bars, ±SD. **P < 0.01 by Student’s t-test. Right panel, mean square relative velocity Δ*v*_RMS_ of pairs of neighboring cells obtained from nuclear tracking. Two cells are considered neighbors if the average distance between the centroids of the respective nuclei is below 14 μm.

The silencing of myosin VI with specific siRNA oligos had no impact on the mean velocity and persistence motility of individual cells in random or confined migration assays (Figure 1B, C, and Video S1,2). Conversely, in wound healing experiments myosin VI depletion caused an impairment of collective migration, consistent with previous findings [31]. The defective collective motion is the result of a reduction in the global velocity of wound closure (Figure S1B and Video S3), likely due to the impairment of cell directionality (Figure S1C). A second siRNA myosin VI oligo showed similar depletion efficiency and almost identical results (Figure S1D, E).

We then analyzed the reawakening of collective migratory properties during unjamming of otherwise solid and static epithelial monolayers. Time-lapse image sequences of control and knock-down (KD) cells were analyzed by particle image velocimetry (PIV) to determine the local prevalent direction of motion and obtain time-resolved coarse-grained velocity fields [14]. As shown in Figure 1D and Video S4-5, myosin VI depletion strongly impacts collective migration by reducing the overall cellular motility, quantified by the root mean square velocity *V*_*RMS*_ (Figure 1E), and, even more strikingly, by severely impairing long-range cell-cell coordination. The degree of mutual alignment of cellular velocities is captured by the polar order parameter *ψ*, which can vary in the range of [0, 1], with *ψ* = 1 corresponding to a perfectly uniform velocity field, and *ψ* ≃ 0 to a randomly oriented velocity field (see Methods for details). While we measure *ψ* ≃ 0.5 for the control monolayer, clearly indicating the presence of directed collective migration, we observed a four-to five-fold decrease in *ψ* for the myosin VI KD condition (Figure 1F). We confirmed the lack of long-range coordination in KD monolayers by calculating the velocity correlation functions *C*_*VV*_(r) and the corresponding correlation lengths *L*_*corr*_ (Figure 1G). *L*_*corr*_,which roughly corresponds to the characteristic linear size of a “pack of coherently migrating cells, displays a striking 20-fold reduction in KD cells. Intriguingly, besides reducing large-scale coordination, myosin VI depletion also enhances small-scale velocity fluctuations, as can be seen by considering the mean squared relative velocity √(<Δ *v*_*RMS*_ >)_2_ of neighboring cells pairs, which displays a significant increase in KD monolayers (Figure 1G). Of note, all these effects are not due to a cell division defect since a similar number of cells were present in fully confluent mock and myosin VI depleted monolayers (Figure S1F) and this collective locomotion is unperturbed by replication inhibition [14].

Altogether, these results showed that the critical role of myosin VI in controlling multicellular streaming-like motility is an emergent property of confluent monolayers.

### Myosin VI coordinates cryptic lamellipodia dynamics in the follower cells

Myosin VI has been reported to be localized at adherens junctions and to interact with the cytoplasmic tail of E-cadherin [32-34]. In MCF10.DCIS.com cells, myosin VI displays a diffuse cytoplasmic pattern in isolated, sparsely-seeded cells but rapidly accumulates at desmosomes when a cluster of two or more cells is established (Figure S2A). In confluent monolayers, myosin VI was also present at the basal level where it accumulated in actin-rich protrusions, called cryptic lamellipodia, which extend underneath the neighboring cells and display a planar polarized distribution [12, 35] (Figure 2A, B). Importantly, by monitoring the dynamics of cells expressing EGFP-LifeAct interspersed with non-fluorescent cells, we found that myosin VI depletion did not impair the formation of cryptic lamellipodia (Figure 2C and Video S6) but severely reduced their velocity (Figure 2D) and persistence (Figure 2E).

**Figure 2.**
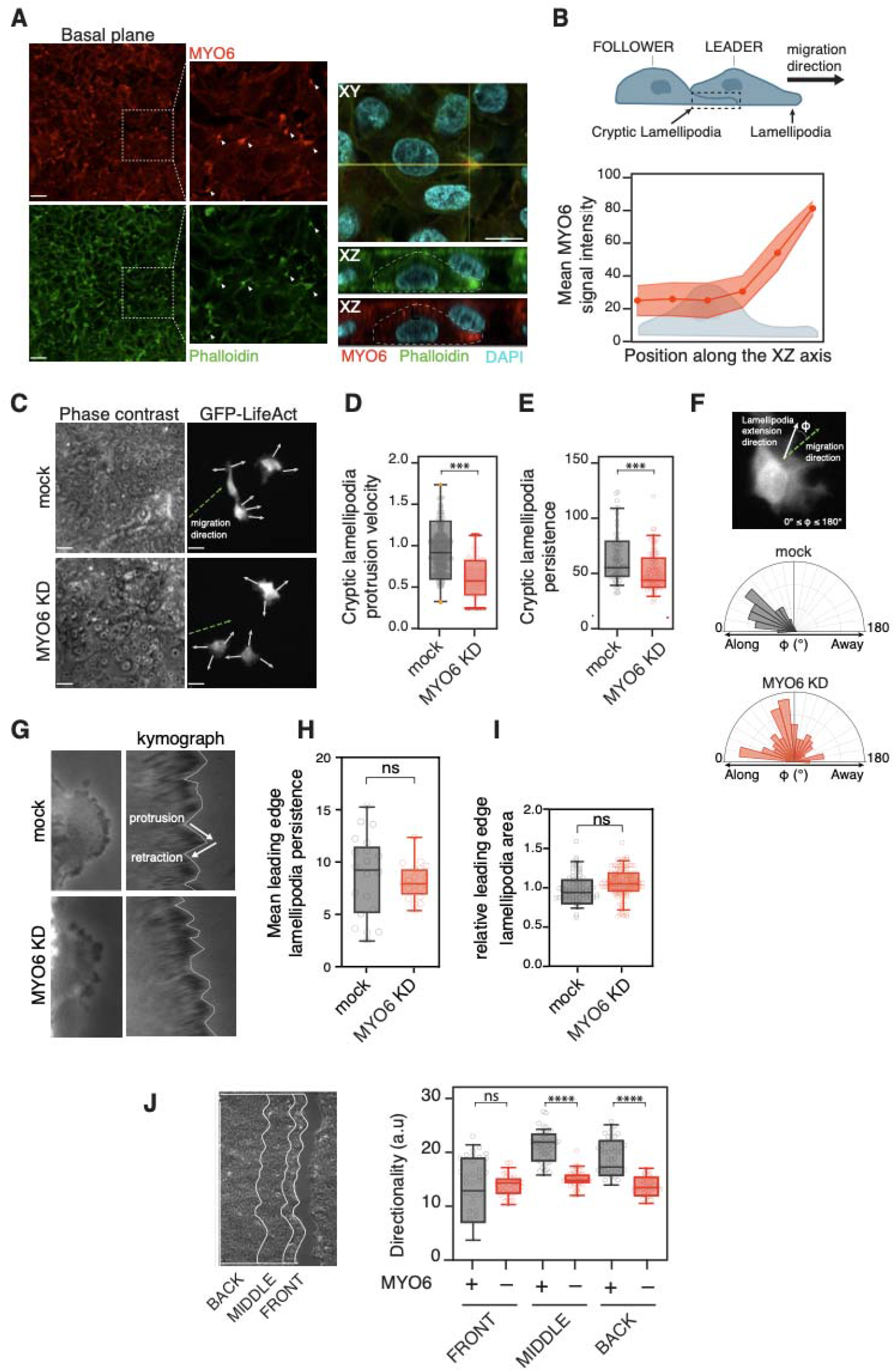
Myosin VI coordinates cryptic lamellipodia dynamics in follower cells. **(A)** Basal plane of a fully confluent DCIS-RAB5A monolayer. Immunofluorescence (IF) analysis was performed as indicated. Arrows in the magnifications indicate the accumulation of myosin VI signal at the cryptic lamellipodia together with phalloidin. Scale bars, 10□μm. Right panel, example of Z-stacks projection of the same cells. Scale bars, 5□μm. **(B)** Top, schematic representation of cryptic lamellipodia structure. Bottom, mean fluorescence signal of myosin VI across single cells from Z-stacks acquisition images. *n*= 60 (15 cells for four independent experiments). Error bars, ±SD. **(C)** Representative phase-contrast and fluorescent images of the streaming assay of DCIS-RAB5A monolayers where GFP-LifeAct-expressing mock or myosin VI depleted cells were interspersed (1:10 ratio). Green arrow, direction of the migrating cell sheet. Scale bars, 15□μm. **(D)** Quantification of cryptic lamellipodia protrusion velocity from the streaming assay described in C. Images of cryptic lamellipodia from time-lapse microscopy were used to monitor the locomotion of the green cells quantified with the ADAPT plug-in of Fiji. Results in the graph are expressed relative to GFP-LifeAct-expressing mock average value. Empty circle represents the mean lamellipodia protrusion velocity of a single cell. *n*≅120 (at least 30 cells/condition for four independent experiments). ***P < 0.001 by Student’s t-test. **(E)** Quantification of cryptic lamellipodia persistence from the streaming assay described in C. Data is plotted as the number of frames in which the lamellipodia is detectable. Empty circle represents the persistence of single lamellipodia *n*≅80 (at least 20 lamellipodia/condition for four independent experiments). ***P < 0.001 by Student’s t-test. **(F)** Image representation of the angle *Φ* between the direction of each cryptic lamellipodia (white arrow) and the direction of the migrating cell sheet (green arrow). *Φ* ∼ 0° indicates that protrusions and collective migration have the same direction; *Φ* > 90° indicates that the directions of protrusions and collective migration are diverging. Right plots: quantification of the orientation angle *Φ*. *n*≅140 (35 lamellipodia/condition for four independent experiments). **(G)** Representative phase-contrast images of leading edge lamellipodia and their relative kymograph from GFP-LifeAct expressing DCIS-RAB5A mock or myosin VI depleted (KD) scratched cell monolayers. Arrows in the kymograph highlight one lamellipodia protrusion and retraction event. **(H)** Quantification of leading edge lamellipodia persistence (expressed in minutes) obtained by kymograph analysis described in G. Empty circle represents the persistence of single leading edge lamellipodia. *n*≅20 (at least 5 lamellipodia/condition for three independent experiments). ns > 0.999 by Student’s t-test. **(I)** Quantification of leading edge lamellipodia mean area by neural network [36] using phase-contrast images of wounded DCIS-RAB5A mock or myosin VI depleted (KD) cell monolayers. Results in the graph are expressed relative to mock average value. Empty circle represents the area occupied by leading edge lamellipodia. *n*≅80 (at least 20 lamellipodia/condition for four independent experiments). ns > 0.999 by Student’s t-test. **(J)** Single cell tracking of the wound healing assay performed on H2B-mCherry expressing DCIS-RAB5A cells, mock or myosin VI depleted (KD). Left panel, representative phase contrast images of wound healing assay. The area to which cells are assigned based on their distance from the wound edge is highlighted with white lines. Right panel, quantification of the different area. The reported directionality is the inverse of the directional change rate parameter obtained with TrackMate. Empty circle represents the mean directionality of the tracks identified in a single timelapse video. *n*≅30 (at least ten videos/condition for three independent experiments). Error bars, ±SEM. ns > 0.999, ****P < 0.0001 by ANOVA test.

One striking feature of flocking-fluid locomotion in DCIS-RAB5A cells is their long-range, persistent, and ballistic motility. This trait is driven by the formation of highly coordinated cryptic lamellipodia oriented in the direction of motion. Myosin VI depletion strongly impaired the alignment of these protrusions along the motility direction of supracellular motility streams (Figure 2F). Notably, myosin VI depletion impaired specifically cryptic lamellipodia in cell monolayers as its depletion did not alter protrusion and persistence of lamellipodia generated in single cells (Figure S2B, C).

Cryptic lamellipodia are typically observed in follower cells during direct cell motility [12]. The streaming motility in monolayers, however, does not allow discerning leader-to-follower topological cell organization. Thus, we analyzed myosin VI activity using a wound healing assay where the migratory phenotype caused by its depletion is evident (Figure S1B, C). For each cell, we measured both its absolute speed and axial component in the direction of the wound by tracking its H2B-mCherry-labelled nucleus. Consistent with the specific role of myosin VI in the followers, the silencing of the protein did not alter lamellipodia extension and persistence in leader cells at the wound front as quantified by kymograph (Figure 2G, H) and neuronal network analyses [36] of the dynamics at the leading edges (Figure 2I). We also employed single-cell tracking analysis to evaluate the movement of cells located away from the wound edge. Strikingly, we found a significant reduction in cell directionality specifically in the follower cells that were a few rows away from the wound edge (Figure 2J).

Collectively, these findings indicate that myosin VI specifically controls the coordinated persistence and dynamics of cryptic lamellipodia in the follower cells, a prerequisite for the collective movement.

### Myosin VI regulates RAC1 GTPase activation at the cryptic lamellipodia

A critical molecular determinant of cryptic lamellipodia is the small GTPases RAC1 whose activity is essential to trigger localized branched actin polymerization at the leading edge of migratory cells. We found that the total amount of active RAC1 was significantly reduced in myosin VI-depleted lysates from confluent cell monolayers (Figure 3A, Figure S3A). Importantly, the loss of myosin VI does not alter the level of active CDC42 (Figure 3B, Figure S3B), nor of RAC1 when lysates are prepared from isolated cells grown in sparse conditions (Figure S3C), reinforcing the notion of a specific function of myosin VI in controlling RAC1 activity in streaming, fluidized monolayers.

**Figure 3.**
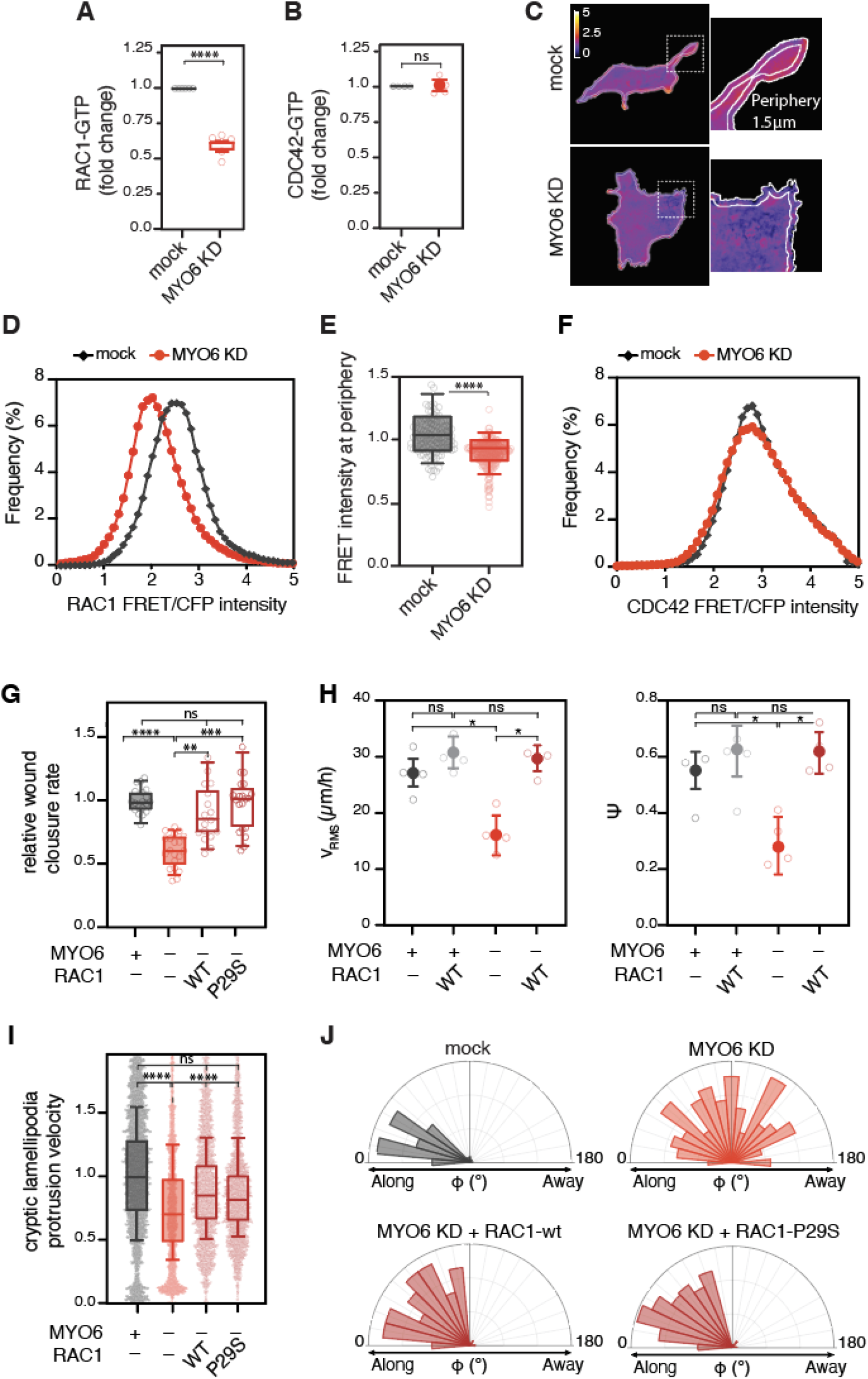
Myosin VI regulates RAC1 GTPase activation at cryptic lamellipodia. **(A)** Quantification of the GST-CRIB assay performed using lysates from DCIS-RAB5A mock or myosin VI depleted (KD) monolayers. The intensity of the RAC1-GTP band was normalized to the total amount of RAC1 present in the lysate. Data are reported as fold change with respect to RAC1-GTP level in the corresponding mock sample for each experiment. *n*=5 independent experiments. Reported values are mean ± SD. ****P < 0.0001 by Student’s t-test. **(B)** As in A, but for CDC42-GTP. Reported values are mean ± SD. ns > 0.999 by Student’s t-test. **(C)** Representative images of RAC1-FRET-biosensor expressing DCIS-RAB5A cells, mock or myosin VI depleted (KD), used for the quantification of the FRET intensity signal at the cell periphery (<1.5 µm from the edge). Cells were seeded in jammed condition mixing FRET biosensor expressing cells in a 1:10 ratio with DCIS-RAB5A cells. The signal was recorded 16 hours after RAB5A induction. **(D)** Representative distribution of the normalized RAC1-FRET intensity signal at the cell periphery of a mock or myosin VI depleted (KD) cell. **(E)** Quantification of the FRET intensity signal at the cell periphery of mock or myosin VI depleted (KD) cells. Results are expressed relative to mock average value. *n*≅120 (at least 30 cells/condition for four independent experiments). ****P < 0.0001 by Student’s t-test. **(F)** Representative distribution of the normalized CDC42-FRET intensity signal at the cell periphery of a mock or myosin VI depleted (KD) cell. **(G)** Wound healing assay of DCIS-RAB5A mock or myosin VI depleted (KD) cells expressing the indicated constructs. Motility was quantified by measuring the area covered over time. The average wound closure speed relative to the mock condition is plotted in the graph. The empty circle represents the mean wound closure velocity quantified for each video. *n*≅20 (at least five videos/condition for four independent experiments). ns > 0.999, **P < 0.01, ***P < 0.001, ****P < 0.0001 by ANOVA test. **(H)** PIV analysis of the streaming assay performed as in Figure 1 E, F for the indicated cell lines. Left panel, root mean squared velocity *V*_*RMS*_, right panel, orientational order parameter *Ψ*. Empty circle, mean of the single experiment calculated from at least five videos/condition for four independent experiments. Error bars ± SD. ns > 0.999, *P < 0.05 by ANOVA test. **(I)** Quantification of cryptic lamellipodia protrusion velocity performed as in Figure 2D for the indicated cell lines. Results in the graph are expressed relative to GFP-LifeAct-expressing mock average value. The empty circle represents the mean lamellipodia protrusion velocity of a single cell. *n*≅140 (at least 45 cells/condition for three independent experiments). ns > 0.999, ****P < 0.0001 by ANOVA test. **(J)** Quantification of the orientation angle *Φ* performed as in Figure 2F. Control mock or myosin VI depleted cells, *n*≅140 (40 lamellipodia/condition for three independent experiments). Cell population expressing RAC1 wild-type (WT) and RAC1-P29S, *n*≅250 (80 lamellipodia/condition for three independent experiments).

Next, we generated a DCIS-RAB5A stable cell line expressing a second-generation RAC1-FRET biosensor [37]. Cells expressing the RAC1-FRET sensor were mosaically seeded in a confluent monolayer composed of non-fluorescent cells to enable the measurement of RAC1 activation with high spatial resolution. In control cells, analysis of multiple protrusions revealed an increase in FRET activity in the proximity of the cell periphery (arbitrarily estimated at 1.5 μm distance from the plasma membrane), consistent with the established role of RAC1 activity in cryptic lamellipodia [38, 39] (Figure 3C-E). Strikingly, the depletion of myosin VI significantly reduced the relative FRET signal of RAC1 activity in protrusions (Figure 3C-E). Importantly, the absence of myosin VI did not alter the activation of CDC42, measured using a specific FRET biosensor [40] (Figure 3F). These results indicate that myosin VI is essential for optimal activation of RAC1 in cryptic lamellipodia that drive coordinated, long-range streaming motility [35].

Finally, we tested whether the constitutive activation of RAC1 was sufficient to rescue the phenotypes induced by myosin VI depletion using different orthogonal approaches. First, we found that the addition of cholinergic agonist and RAC1 activator carbachol [41, 42] partially rescued the wound closure defect observed upon myosin VI depletion (Figure S3D). Next, we expressed RAC1 wild-type (WT-RAC1) or its fast cycling RAC1-P29S variant [43] in EGFP-LifeAct-expressing cells after silencing of myosin VI. Both WT-RAC1 and RAC1-P29S were sufficient to rescue the wound closure defect (Figure 3G and Video S7) as well as the streaming defect observed in myosin VI depleted cells upon RAB5A-induced unjamming. Indeed, the ectopic expression of RAC1 in the cell monolayer was sufficient to rescue the loss of cell coordination measured by spatial velocity correlation lengths and root mean square velocity (Figure 3H and Video S8). Finally, we tested the dynamics of cryptic lamellipodia in the flocking, confluent monolayers. The expression of either WT-RAC1 or RAC1-P29S effectively restored the defective cryptic lamellipodia velocity (Figure 3I) and their directional orientation (Figure 3J) caused by myosin VI depletion.

Altogether, these results demonstrate that myosin VI regulates the localized activation of RAC1 specifically at cryptic lamellipodia protrusions to promote coordinated and collective cell migration and tissue fluidification.

### Myosin VI controls RAC1 activation in cryptic lamellipodia by recruiting DOCK7

To gain insight into the molecular mechanisms underlying myosin VI regulation of RAC1 activity, we took advantage of the myosin VI interactome, which we identified in a previous study [25]. Among the myosin VI interactors, we focused on the human dedicator of cytokinesis DOCK7 [31, 44, 45], a dual guanine nucleotide exchange factor (GEF) for RAC1 and CDC42 GTPase [46]. We first confirmed that myosin VI and DOCK7 co-immunoprecipitated in DCIS-RAB5A cells (Figure 4A). Next, we tested the functional involvement of DOCK7 in the regulation of RAC1 activity in cryptic lamellipodia in our model of flocking fluid motility in monolayers.

**Figure 4.**
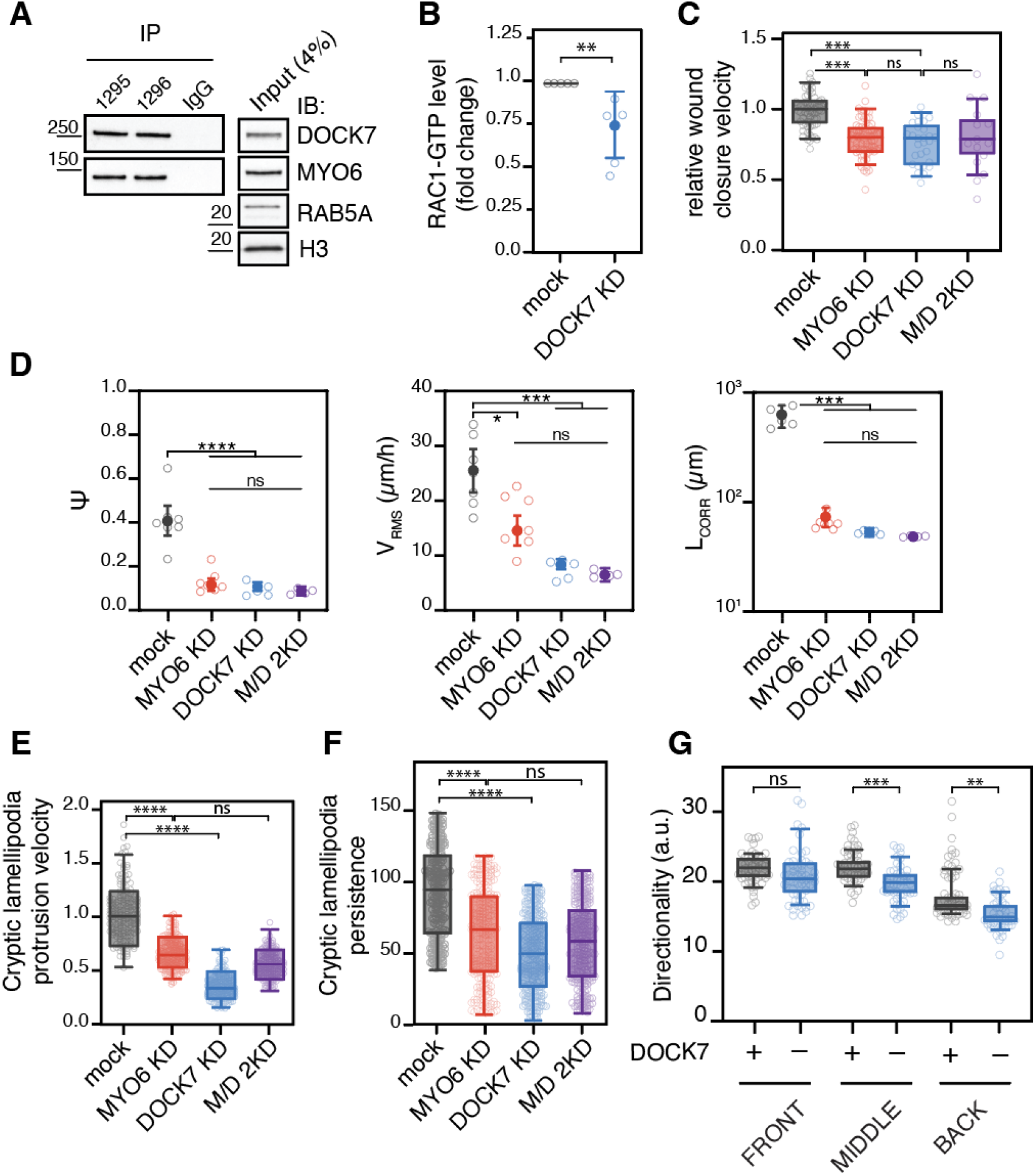
A MYO6-DOCK7 axis activates RAC1 in a planar polarized fashion. **(A)** Immunoprecipitation (IP) analysis of DCIS-RAB5A cells with two antibodies against myosin VI (1295 and 1296) and rabbit IgG as a negative control. IB as indicated. **(B)** Quantification of the GST-CRIB assay performed using lysates from DCIS-RAB5A mock or DOCK7 depleted (KD) monolayers. The intensity of the RAC1-GTP band was normalized to the total amount of RAC1 present in the lysate. Data are reported as fold change with respect to RAC1-GTP level in the corresponding mock sample for each experiment. *n*=5 independent experiments. Reported values are mean ± SD. **P < 0.01 by Student’s t-test. **(C)** Wound healing assay of DCIS-RAB5A mock, MYO6 and DOCK7 depleted cells, singularly or in combination (M/D 2KD) cells. The motility rate was quantified by measuring the wound-closure area covered over time. The average wound closure speed relative to the mock condition is plotted in the graph. The empty circle represents the mean wound closure velocity quantified for each video. *n*≅20 (at least five videos/condition for four independent experiments). ns > 0.999, ***P < 0.001 by ANOVA test. **(D)** PIV analysis of the streaming of DCIS-RAB5A mock, MYO6, and DOCK7 depleted singularly or in combination (M/D 2KD) cells. From left to right: orientational order parameter *Ψ*, root mean squared velocity *V*_*RMS*_, and correlation length *L*_*corr*_. Empty circle, mean of each experiment calculated from at least five videos/condition. The data reported are from at least six independent experiments. Error bars ± SD. ns > 0.999, *P < 0.05, ***P < 0.001, ****P < 0.0001 by ANOVA test. **(E)** Quantification of cryptic lamellipodia protrusion velocity performed as in Figure 2D for the indicated cell lines. Protrusion velocity data are plotted relative to the mock condition. The empty circle represents the mean lamellipodia protrusion velocity of a single cell. *n*≅160 (at least 40 cells/condition for four independent experiments). **(F)** Quantification of cryptic lamellipodia protrusion persistence, performed as in Figure 2F for the indicated cell lines. Data are plotted as the number of frames in which the lamellipodia is detectable. The empty circle represents the persistence of single lamellipodia. *n*≅240 (at least 60 lamellipodia/condition for four independent experiments). ns > 0.999, ****P < 0.0001 by ANOVA test. **(G)** Quantification of the wound healing assay upon DOCK7 depletion. Cell tracks obtained following mCherry nuclei with the Fiji TrackMate plugin were assigned to the different areas, as described in Figure 2G. The plotted directionality is the inverse of the directional change rate parameter obtained with TrackMate analysis. The empty circle represents the mean directionality of the tracks identified in a single timelapse video. *n*≅30 (at least ten videos/condition for three independent experiments). Error bars, ±SEM. ns > 0.999, ****P < 0.0001 by ANOVA test.

Biochemically, DOCK7 depletion significantly reduced RAC1-GTP levels (Figure 4B). Functionally, DOCK7 KD cells showed impaired collective migration in both wound healing (Figure 4C and Video S9) and streaming assay (Figure 4D and Video S10) to the same extent as in myosin VI KD cells. Intriguingly, the concomitant silencing of both DOCK7 and myosin VI did not worsen the migratory defects, suggesting that DOCK7 is likely the main effector of myosin VI activity in this context (Figure 4C, D). Results were confirmed by a second siRNA DOCK7 oligo (Figure S4A, B).

Next, we tested whether DOCK7-dependent impairment of flocking monolayer migration was due to a lack of oriented and persistent cryptic lamellipodia. By exploiting EGFP-LifeAct mosaic cells in confluent monolayers, we discovered that both protrusion velocity and persistence of cryptic lamellipodia are similarly impaired after either individual or concomitant depletion of DOCK7 and myosin VI (M/D 2KD) (Figure 4E, F). We then used cell segmentation and tracking of H2B-mCherry-labelled monolayer cells in a wound healing assay to analyse specifically the migration of the leader or the follower cells. Consistent with data obtained in myosin VI KD cells (Figure 2G-I), the silencing of DOCK7 did not affect the lamellipodia dynamics of the cell leading edge at the wound front (Figure S4C, D), while kymograph-based quantification of follower cells showed a significant reduction in directionality (Figure 4G).

Prompted by these results, we reasoned that myosin VI may be required to localize DOCK7 and spatially restrict its activity toward RAC1. To test this hypothesis, we first examined DOCK7 localization in our cell system. To overcome the lack of reliable antibodies, we generated a population of EGFP-DOCK7 cells that express low physiological levels of the protein (Figure S4E). A confocal analysis demonstrated that in a confluent monolayer DOCK7 co-localized with myosin VI at apical cell-cell junctions (Figure 5A) and accumulated in cryptic lamellipodia extending basally onto the cell substrate (Figure 5B, C). This localization requires myosin VI as, upon myosin VI depletion, DOCK7 became diffusely distributed throughout the cytoplasm (Figure 5D) and was no longer enriched at actin-rich protrusion tips (Figure 5E, F).

**Figure 5.**
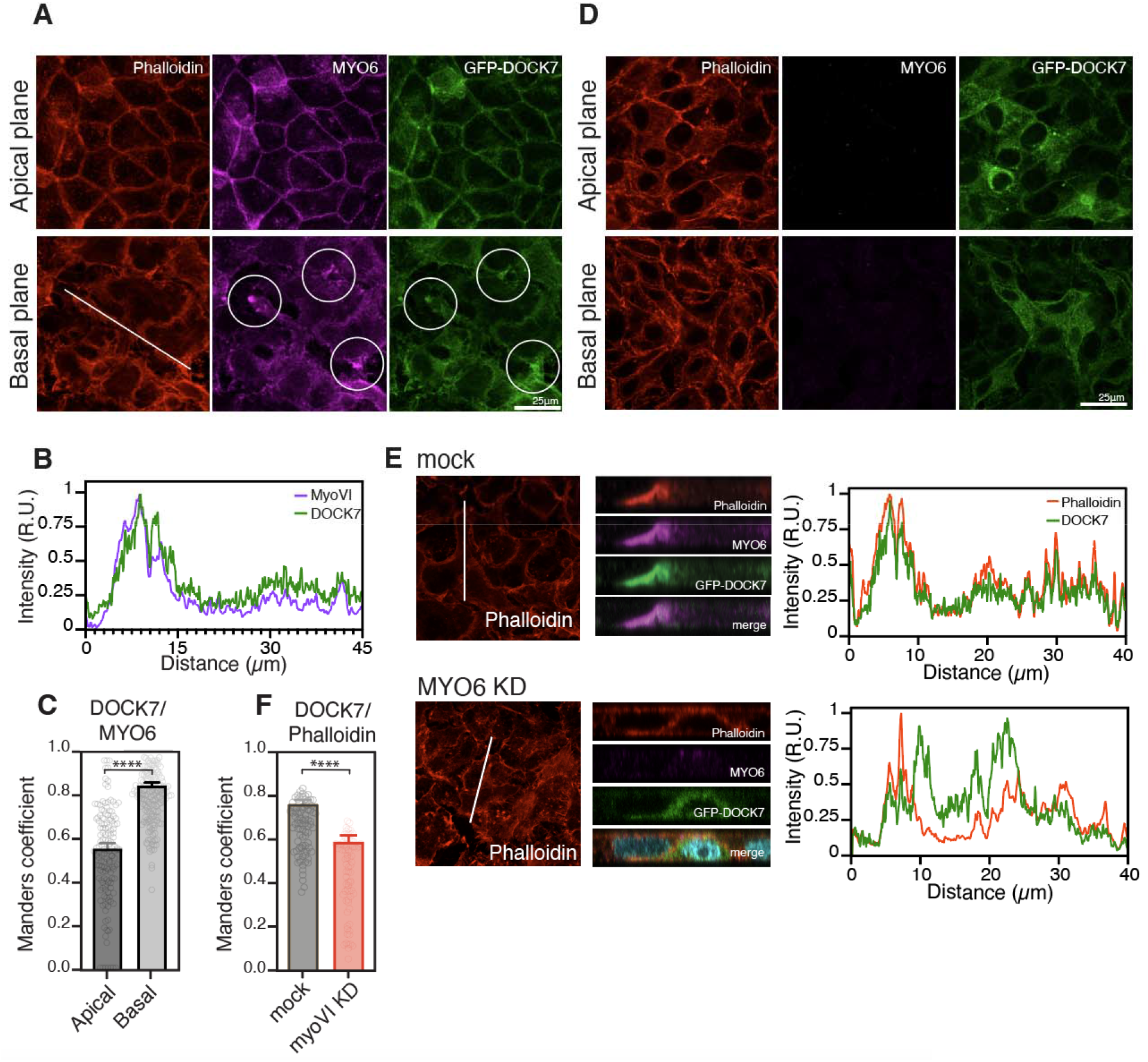
Myosin VI promotes local RAC1 activation by recruiting DOCK7 to cryptic lamellipodia. **(A)** IF analysis of GFP-DOCK7 expressing DCIS-RAB5A cells seeded in jammed condition to visualize lamellipodia-like structures. Purple, myosin VI; red, phalloidin. Middle and basal planes are shown. Scale bars: 25µm **(B)** Fluorescence intensity profiles show GFP-DOCK7 and MYO6 fluorescence distribution across the white line shown in A (x-axis). The fluorescence intensities are reported on the y-axis. The overlay of the two signals indicates colocalization. **(C)** Quantification of the colocalization of GFP-DOCK7 and myosin VI shown in A, using Manders’ coefficient. *n*=150 in at least 35 fields of view from four independent experiments. Error bars, ±SEM. ****P < 0.0001 by Student’s t-test. **(D)** IF analysis of GFP-DOCK7 expressing DCIS-RAB5A cells depleted of myosin VI and seeded in jammed condition, as in A. Purple, myosin VI; red, phalloidin. Middle and basal planes are shown. **(E)** Z-stacks acquisition of the cell lines described in A and D. Right panel, fluorescence intensity profiles showing the distribution of GFP-DOCK 7 and phalloidin fluorescence across the white line (x-axis). **(F)** Quantification of the colocalization of GFP-DOCK7 and phalloidin in XZ images shown in E, using Manders’ coefficient. *n*=80 in at least 20 fields of view from four independent experiments. Error bars, ±SEM. ****P < 0.0001 by Student’s t-test.

Collectively, these data indicate that a myosin VI-DOCK7-RAC1 axis controls cryptic lamellipodia protrusions, which are, in turn, required for collective flocking locomotion.

### Myosin VI directly interacts with the DOCK7 DHR2 domain

A functional interaction between myosin VI and DOCK7 has been previously reported in the neuronal context [47] as well as HeLa [44] and HEK293T cells [31], but the molecular basis of this interaction has not been fully explored. Structurally, both proteins are composed of several, distinct domains that we investigated to map the critical surface of interaction (Figure S5A). First, we confirmed that the myosin VI binding surface resides in the DHR2 domain of DOCK7 [45], as the removal of this GEF catalytic domain in the context of the full-length protein was sufficient to abrogate binding to the myosin VI tail (Figure 6A). To identify the minimal binding region within myosin VI, we performed a pull-down experiment with different myosin VI tail constructs. Both the cargo-binding domain (CBD, [44]) and the MYO6 ubiquitin-binding (MyUb, [48]) isolated domains bound DOCK7, although with reduced efficiency as compared to the MyUb-CBD tail of myosin VI (Figure S5B). Importantly, using bacterially expressed and purified fragments, we showed that the interaction between the DHR2 domain and the MyUb-CBD domain is direct (Figure 6B).

**Figure 6.**
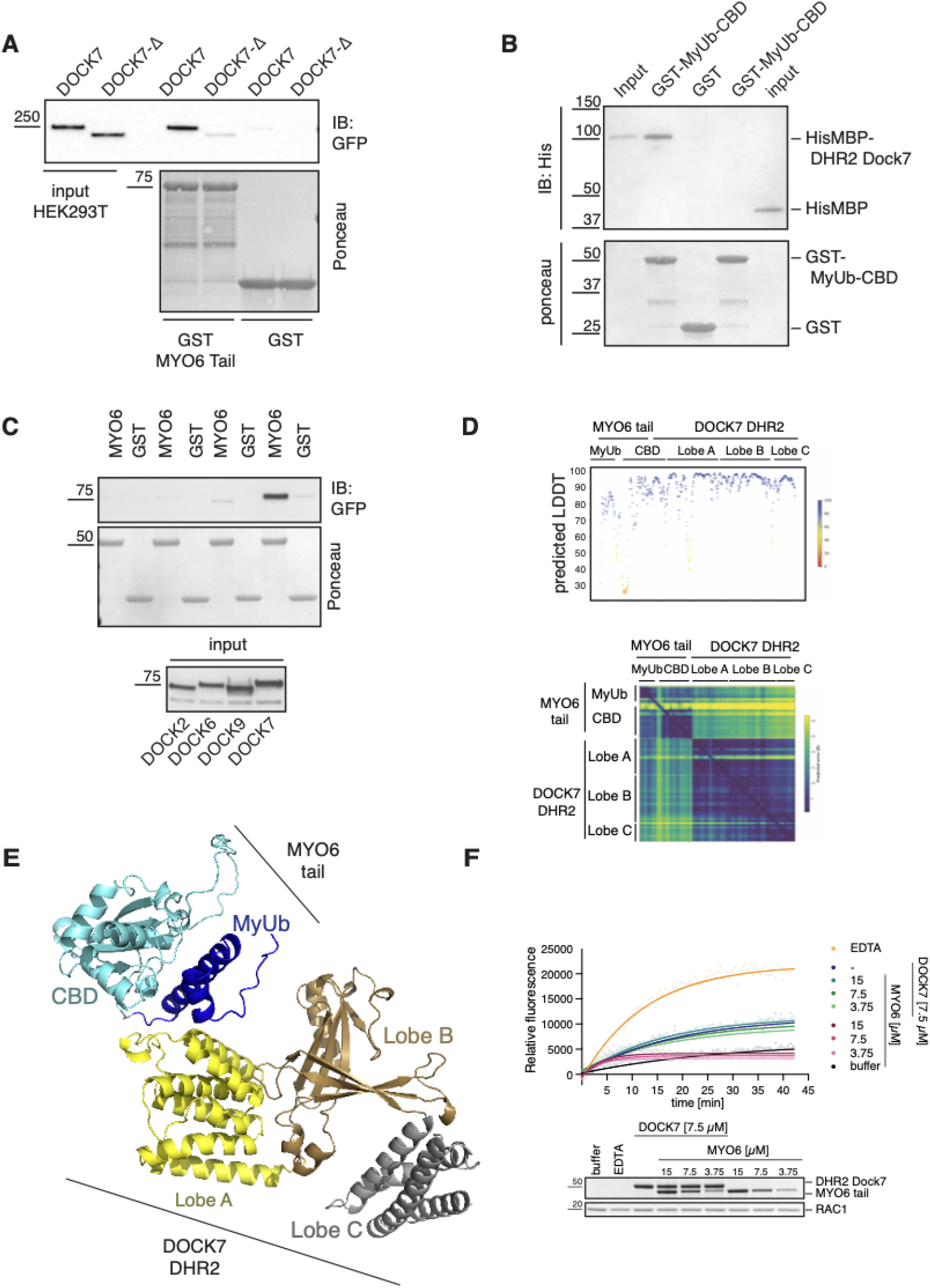
Myosin VI specifically and directly interacts with the Lobe A of the DHR2 domain of DOCK7. **(A)** GST pull-down assay using myosin VI tail and lysates from HEK293T cells transfected with full-length GFP-DOCK7 or its DHR2 deleted mutant, GFP-DOCK7ΔDHR2 (DOCK7Δ). IB as indicated. Ponceau was used to show equal loading. **(B)** Pull-down assay using HisMBP-DHR2 domain of DOCK7 and MyUb-CBD construct of myosin VI produced and purified from bacteria. IB as indicated. Ponceau was used to show equal loading. **(C)** GST pull-down assay using MyUb-CBD of myosin VI (spanning amino acids 1080-1295) and lysate from HEK293T cells transfected with GFP-DHR2 domain of the indicated DOCK proteins. IB as indicated. Ponceau was used to show equal loading. **(D)** Ribbon diagram of the top-scoring AlphaFold2-Multimer model of the MYO6:DOCK7 interaction obtained by inputting protein sequences for the myosin VI tail, spanning residues G1048 – K1262, and DOCK7 residues of the DHR2 domain spanning P775 – P1196. **(E)** Confidence scores per residue generated by AlphaFold2-Multimer for the predicted fold of domains (upper panel) and the residue-to-residue distance (bottom panel). LDDT, local distance difference test. **(F)** Representative RAC1 GEF activity assay using 7.5 µM of the DHR2 domain of DOCK7 and the indicated concentration of the MyUb-CBD myosin VI. Bottom, Coomassie gel showing the correct loading of the indicated proteins.

The DOCK family consists of 11 structurally conserved proteins that serve as atypical RHO GEFs and are differentially expressed in tissues [49, 50]. We tested the ability of the MyUb-CBD tail to bind the DHR2 domain of a few prototypes of the family, including DOCK2, DOCK6 and DOCK9 [51]. Surprisingly, binding was detected only for the DHR2 domain of DOCK7 (Figure 6C).

This result prompted us to further analyze the interaction surface. Despite low sequence homology among the DOCK family members, the DHR2 domains are well-conserved and adopt a similar fold that is characterized by three lobes, A–C. Of them, Lobes B and C are endowed with the GTPase binding and GEF activity, whereas Lobe A seems to be involved in homodimerization, at least in a few DOCK proteins [49, 51]. By using DHR2 protein fragments, we showed that Lobe A is, indeed, required for DOCK7 dimerization, but is also critical for myosin VI interaction (Figure S5B). Lobe A is not present in the recombinant DHR2 constructs used to generate the structural data for DOCK7 (pdb6AJ4). Therefore, we used AlphaFold2-Multimer [52, 53] to predict the DOCK7 DHR2 structure and possible myosin VI interaction surfaces. The DOCK7 DHR2 and myosin VI MyUb and CBD domains were predicted with high confidence scores except for a linker sequence between the two myosin VI domains that most likely is flexible (Figure 6D, upper panel). Best model prediction indicated that the DOCK7 Lobe A domain is in contact with the CBD and MyUb domains of myosin VI, with the MyUb and CBD domains also interacting with each other (Figure 6E). This model showed high confidence for the residue-to-residue distance (Figure 6D, bottom panel) and is fully consistent with the finding that loss of either the MyUb or CBD weakens the interaction with DOCK7 (Figure S5B).

A hypothesis motivated by our structure-function analysis is that the binding of myosin VI to Lobe A may influence the GEF activity of DOCK7. Indeed, DOCK7 showed poor activity on RAC1 compared to DOCK2 (Figure S5C) as previously reported [46], strongly suggesting a possible allosteric missing partner. We then used a suboptimal concentration of DOCK7 and titrated in increasing amounts of the MyUb-CBD tail (Figure 6F), analyzing activity by the GEF assay. Even under these conditions, however, we failed to detect any effect of the MyUb-CBD fragment on the GEF activity of DOCK. Thus, we conclude that, while critical for DOCK7 localization, myosin VI does not appear to influence DOCK7 GEF activity toward RAC1, at least in this simplified *in vitro* experiment.

### Myosin VI overexpression is exploited by infiltrating breast cancer cells

To assess the clinical relevance of our findings, we investigated the expression profile of myosin VI in human breast cancer, by analyzing RNA-seq data of The Cancer Genome Atlas (TCGA) breast tumors dataset (BRCA). We assessed a total of 981 samples with complete clinical and pathological information including molecular subtyping [54] and we focused our attention on the myosin VI short isoform, as we previously demonstrated that this isoform is selectively required for cancer cell migration and for DOCK7 binding [31]. As shown in Figure 7A, the expression of myosin VI short isoform is significantly higher in the basal-like subtype when compared to all other cancer subtypes and normal breast tissues. Notably, this subtype comprises 15-20% of all breast tumors and is assimilated to triple-negative (TN) highly metastatic breast cancers to which the MCF10.DCIS.com cell line belongs. To confirm this result at the protein level, we analyzed a panel of human ductal in situ carcinoma (DCIS) and invasive ductal carcinoma (IDC) tissue sections by immunohistochemistry. While a diffuse and weak expression of myosin VI characterized most DCIS samples, the staining intensity was significantly higher in IDC, particularly in the infiltrating components (Figure 7B) as quantified by software analysis on digital slide scans (Figure 7C). Thus, breast carcinoma selectively increase myosin VI expression during the progression from DCIS to IDC.

**Figure 7.**
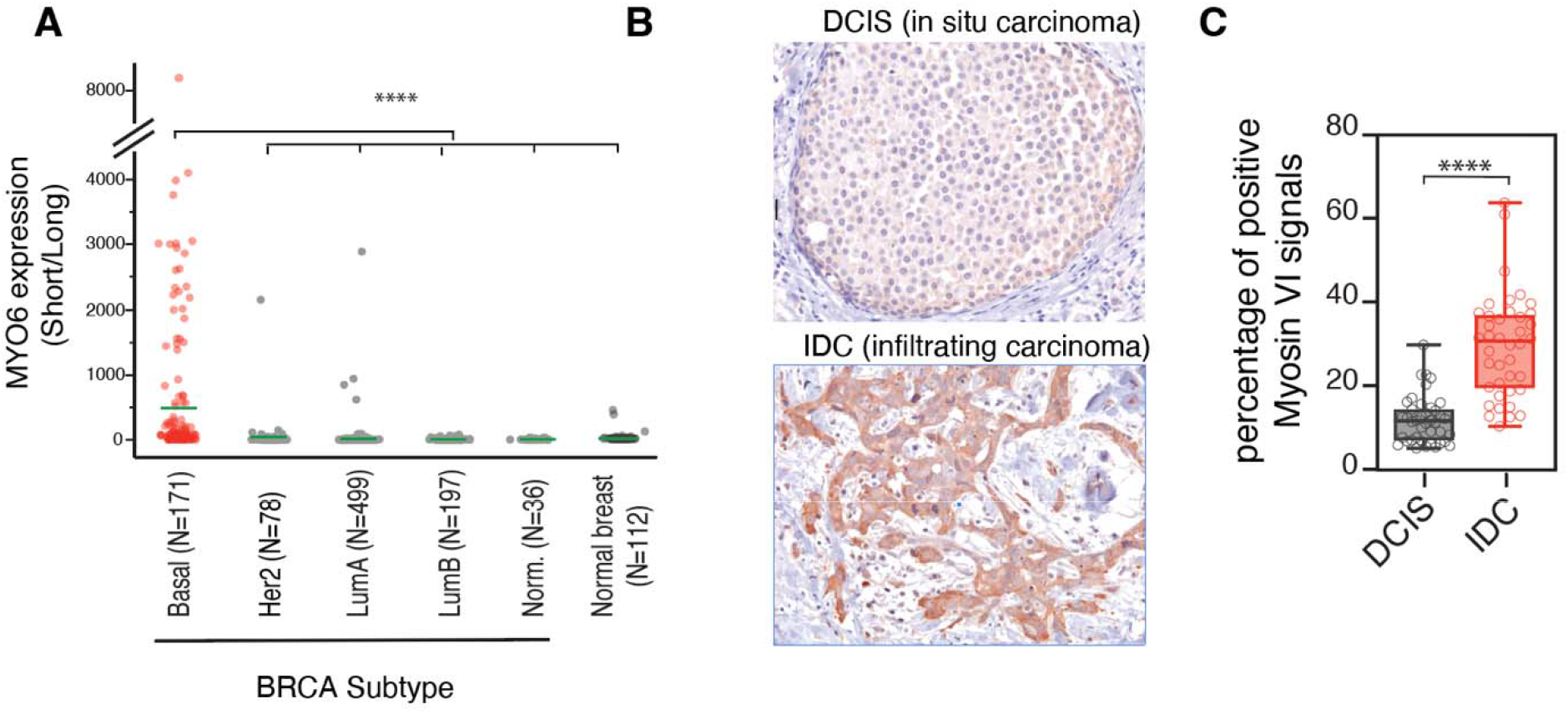
MYO6 is significantly overexpressed in highly metastatic breast cancers. **(A)** MYO6 short versus long isoform expression expressed as ratio of RPKM for the indicated BRCA subtypes. Average expression level is shown by green lines. Myosin VI short isoform was significantly overexpressed in BRCA basal-like subtype when compared to all other subtypes and normal breast tissues. Statistical significance was analyzed by one-way (Chi-Square approximation) Wilcoxon Test. ***P < 0.0001. **(B)** Representative immunohistochemistry (IHC) images of myosin VI in DCIS and IDC tumor sections. Original magnification, x200. Scale bar, 100 µm. **(C)** Quantitative analyses of myosin VI IHC images shown in B. n=40 (5 images of 8 cases/cancer type). ****P < 0.0001 by Student’s t-test.

## DISCUSSION

During carcinoma dissemination, cellular rearrangements are fostered by a solid-to-liquid transition known as unjamming through partially identified molecular mechanisms. We found, here that myosin VI is essential to support this tissue-level phase transition, as its depletion severely reduces cell coordination and impairs cell migration persistence and directionality. Molecularly, we identified DOCK7, a GEF for RAC1, as the critical and direct myosin VI interactor. Myosin VI is essential to restrict DOCK7 at the cryptic lamellipodia to locally activate RAC1 and to promote coordinated movement of the follower cells. This regulation may likely aid the follower cells to chase and coordinate their motion with the leaders, as recently suggested by several studies [55, 56] thereby enabling maintainance of monolayer compactness during collective motion. Our results also highlight the role exerted by RAC1 in the follower cells and show that these cells are not simply hitchhikers or passive passengers, but rather actively contribute to promoting collective cell migration.

Our study has striking similarities with recent discoveries obtained in Drosophila by the group of Denise Montell who uncovered the role of a Scrib/Cdep/Rac pathway as being essential for follower-cell movement and cluster cohesion in border cell migration [55]. In this study and context, Cdep was identified as the Rac GEF, whereas Scrib, Dlg, and Lgl aid in localizing Cdep basolaterally to activate Rac in followers. Intriguingly, Drosophila was the system first employed to demonstrate the critical role of myosin VI in collective motion as its depletion severely affects border cell migration [27]. Whether myosin VI does so by perturbing RAC1 activity in the follower cells in conjunction with or alternatively to Scrib and Cdep has not been addressed. Likely, more than a RacGEF is required in follower cells and little is known about the DOCK7 ortholog in Drosophila, namely Zir. Thus, it will be exciting to re-evaluate the role of myosin VI and Zir activity in border cell dynamics in the light of our current finding.

It must be noted that border cells display not only a leader-to-follower topological organization but also an apicobasal polarity during their motion, consistent with their prototypical epithelial nature. Conversely, breast carcinoma MCF10.DCIS.com cells nearly completely lose their apico-basal polarity while they retain a number of features of normal epithelial tissues including a planar polarized organization. As the molecular determinants of these polarity programs are distinct, it is conceivable that myosin VI might be more critical when a planar polarity arrangement is needed and established, but is dispensable during apico-basal organization. This specific role is particularly attractive considering the selective role exerted by the alternatively spliced myosin VI isoforms [25, 31]. Indeed, fully polarized epithelia selectively express myosin VI long, which is impaired in DOCK7 binding [31] and is critical for clathrin-mediated endocytosis at the apical surface [57].

Finally, the identification of a specific MYO6-DOCK7-RAC1 axis critical for collective cell migration offers new potential possibilities for clinical treatment, particularly in the case of the more aggressive basal-like breast cancers for which we have limited therapeutic options. Indeed, our previous studies [31] and current bioinformatic analyses (Figure 7A) indicate that the myosin VI short isoform is selectively overexpressed in basal-like breast cancers. While the direct inhibition of myosin VI may have unwanted deleterious effects in normal tissues [58], the interaction surface with DOCK7 represents a promising target to explore in future drug discovery studies.

## Acknowledgments

We thank Alessandro Poli for the support in the single-cell confined migration assay. We are grateful to all members of the IFOM imaging facility, in particular, Francesca Casagrande, Serena Magni and Zeno Lavagnino for support in image acquisition and analysis, and Maria Grazia Totaro for support in cell sorting. This work was supported by the Associazione Italiana per Ricerca sul Cancro (Investigator grant 2017□19875 to S.P; 2019-18621 and 5Xmille #22759 to G.S; My First AIRC Grant 2018-22083 to F.G.), and the Seal of Excellence (SoE SEED2020 to S.P. and F.G.) and the Intramural Research Program through the Center for Cancer Research, National Cancer Institute, National Institutes of Health (ZIA BC011627 to K.J.W.). AlphaFold2-Multimer was ran using computational resources of the High-Performance Computing Biowulf cluster of the NIH (http://hpc.nih.gov). Luca Menin’s work was supported by a fellowship of the Associazione Italiana per Ricerca sul Cancro.

## Authors Contribution

Conceptualization, L.M., G.S, and S.P.; investigation, L.M. and J.W.; methodology, L.M., J.W., S.V., El.M., V.C., P.M., A.P., E.F., F.B., C.T. and K.W.; formal analysis, L.M., J.W., S.V.,

Em.M., F.B. and K.W.; supervision, F.G., G.S. and S.P; data curation, L.M. and S.P; writing— original draft, L.M. and S.P.; writing—review & editing, G.S. and S.P.; funding acquisition, S.P.

