## Supplemental information for "A planar-polarized MYO6-DOCK7-RAC1 axis promotes tissue fluidification in mammary epithelia"

**STAR METHODS**

**Constructs and Reagents**

GTP-CRIB plasmid [1] and GST-myosin VI (human isoform2, NP_001287828.1) constructs were previously described [2]. pLenti-RAC1-2G FRET Biosensor and pLenti-Cdc42-2G FRET Biosensor were obtained from Addgene (#66111 and #68813, respectively). DHR2 domain of DOCK2, DOCK6 and DOCK9 GEF proteins were amplified by PCR (primers listed in the “Key resources table”) using cDNA from Caco2 cells, cloned into the expression vector pEGFP-C1 and sequence-verified. pEGFP-DOCK7 was amplified by PCR from pCA-FLAG-DOCK7-FL[1] and cloned into the pEGFP-C1 expression vector. pSLIK-EGFP-DOCK7 and pSLIK-RAC1 wild-type (WT) and P29S constructs were generated starting from previous constructs (pCDNA3-RAC1 [2] and pEGFP-DOCK7) with Gateway™ LR Clonase™ II Enzyme mix (Thermo Fisher Scientific) by subcloning their respective PCR products into a pENTR vector, followed by recombination into the pSLIK-HYGRO empty vector.

All the other truncated constructs were engineered by site‐directed mutagenesis or recombinant PCR and sequence‐verified. Briefly, the DHR2 domain of DOCK7 (residues 1597-2072) was amplified and cloned into the pET28a, pET43 and pEGFP-C1 expression vectors. RAC1 (residues 1-177) was amplified and cloned into the pGEX-6P-1 expression vector. DHR2-ΔLobeA of DOCK7 (residues 1795-2072) was amplified and cloned into the pGEX-6P-1 and pEGFP-C1 expression vectors. Details are available upon request.

**Cell lines and transfection procedures**

The MCF10A-DCIS.com cell line and MCF10A-DCIS.com expressing RAB5A (DCIS-RAB5A) and derivatives stably expressing GFP-LifeAct or mCherry-H2B were previously described [3]. Cells were grown at 37 °C in humidified atmosphere with 5 % CO_2_ in Dulbecco’s Modified Eagle Medium (DMEM): Nutrient Mixture F-12 (DMEM/F12) medium (Invitrogen) supplemented with 5 % horse serum, 0.5 mg/mL hydrocortisone, 10 μg/mL insulin, and 20 ng/mL EGF. Phoenix-AMPHO cells (American Type Culture Collection, CRL-3213) were used as the packaging cell line for the generation of retroviral particles and cultured as recommended by the supplier. HEK293T cells were obtained from the BBCF-Biological Bank and Cell factory, INT, Milan, grown in DMEM supplemented with 10 % fetal bovine serum and 2 mM l-glutamine and used as the packaging line for lentiviral vectors.

DCIS-RAB5A cells and derivatives were infected with pSLIK-EV (empty vector, CTR), pSLIK-EGFP-DOCK7-FL or pSLIK-RAC1 WT and P29S and selected with hygromycin to obtain stable inducible cell lines. In the case of DCIS-RAB5A expressing pSLIK-EGFP-DOCK7-FL, cells were FACS sorted with Beckman Coulter MoFlo Astrios to obtain a homogeneous population with low expression of GFP-DOCK7 in order to determine its localization. In these cell lines, doxycycline induction promotes the expression of RAB5A and DOCK7 or RAB5A and RAC1 WT/P29S. All cell lines were authenticated by cell fingerprinting and tested for mycoplasma contamination.

FRET-based analysis of RAC1 and CDC42 activation was performed infecting DCIS-RAB5A cells with pLent-RAC1-2G [4] or pLenti-CDC42-2G [5] second generation FRET biosensors, followed by selection with puromycin to obtain stable inducible cell lines.

Transient knock-down (KD) of myosin VI or DOCK7 was performed using Stealth siRNA oligonucleotides (listed in the “Key resources table”) from Thermo Fischer Scientific (Waltham, MA, USA). Cells were transfected twice using RNAiMax (Invitrogen), first in suspension and the following day in adhesion. Based on immunoblot analyses, cells were considered myosin VI or DOCK7-depleted three days after the first transfection. Two siRNA oligonucleotides/gene were used with comparable results.

Expression of myosin VI isoforms in DCIS-RAB5A cells was assessed by RT-PCR. RNA was isolated from cells grown at different confluencies using Maxwell® RSC Instrument and Maxwell® RSC simplyRNA Cells Kit. Retro-transcription was performed with QuantiTect Reverse Transcription Kit (Qiagen). The cDNA obtained was amplified by PCR using primers flanking the spliced regions (listed in the “Key resources table”) as previously described [6].

**Antibodies**

The following antibodies were used at the indicated dilutions: anti-His (mouse, 1:1000, Cell Signaling), anti-RAC1 (mouse, 1:1000, BD), anti-E-cad (mouse, 1:200, BD-610181), anti-Desmoplakin (rabbit, 1:1000, NW6; kindly provided by Kathleen Green and Lisa Godsel), anti-myosin VI (rabbit, homemade [6], 1:5000, Eurogentec-1296), anti-GAPDH (mouse, 1:2000, Santa Cruz-32233), anti-DOCK7 (mouse, 1:1000, Santa Cruz-398888), anti-RAB5A (mouse, 1:1000, Santa Cruz-166600), anti-CDC42 (rabbit, 1:1000, Santa Cruz-2462), and anti-GFP (rabbit, 1:5000, Sigma-G1544).

**Single cell migration assay**

Random single cell migration experiment was performed as previously described [7]. Briefly, DCIS-RAB5A mCherry-H2B cells were transfected twice with siRNA oligos and seeded in sparse cell growth conditions in six-well plates (5x10^4^ cells/well) in complete medium. RAB5A expression was induced 16 hours before the experiment was initiated by adding fresh complete media supplemented with 2.5 μg/mL doxycycline to the cells. Cell migration was monitored by time-lapse microscopy. At the time of recording, fresh media containing EGF was added to the cells. The assay was performed using an environmental microscope incubator set to 37 °C and 5 % CO2 perfusion. An Olympus ScanR inverted microscope with 10× objective was used to acquire images every 5 minutes over a 24-hour period. Tracking of cell nuclei and motility analysis were performed as described below.

For confined migration experiment, we employed fibronectin-coated micro-patterned lines of 10 μm diameter created through photolithography [8]. Briefly, glass coverslips were activated with plasma cleaner (Harrick Plasma), followed by coating with PLL-g-PEG (Surface Solutions GmbH, 0.1 mg/mL). After washing with phosphate-buffered saline (PBS), the surface was illuminated with UV light (UVO Cleaner, Jelight) using chromium photo-masks (JD-Photodata). The coverslips were then incubated with fibronectin (25 μg/ml), and 10^4^ DCIS-RAB5A mCherry-H2B transfected cells were seeded over-night with 2.5 μg/mL doxycycline prior to analysis. Images were acquired every 5 minutes for 24 hours using a humidity- and temperature-controlled inverted wide-field ScanR microscope. Tracking of cell nuclei and motility analysis were performed as previously described using a developed C++ software with the OpenCV [http://opencv.willowgarage.com/wiki/] and the GSL [http://www.gnu.org/software/gsl/] libraries. The migration analysis was performed by the C++ software coupled with R [www.R-project.org].

**Wound healing assays**

Assays were performed as previously described [7]. Briefly, cells transfected in suspension were seeded at confluency in six-well plates (1.5x10^6^ cells/well) in complete medium and transfected again the following day. Three days after seeding, a uniform monolayer is formed. RAB5A, DOCK7 and RAC1 WT/P29S expression was induced 16 hours before the experiment was initiated by adding fresh complete media supplemented with 2.5 μg/mL doxycycline to the cells. The cell monolayer was scratched with a pipette tip and carefully washed with PBS to remove floating cells and create a cell-free wound area. The closure of the wound was monitored by time-lapse microscopy. At the time of recording, fresh media containing EGF was added to the cells. The assay was performed using an environmental microscope incubator set to 37 °C and 5 % CO_2_ perfusion. An Olympus ScanR inverted microscope with 10× objective was used to acquire images every 5 min over a 24-hour period. The percentage of area covered by cells (area coverage %) over time and wound-front speed were calculated using a custom Fiji and Matlab code. The area covered over time was fitted with a straight line whose slope was used to estimate the velocity of wound closure.

**Measurement of the cellular velocities and trajectories**

To measure velocity and trajectories of the cells at various distance from the edge of the wound we used Fiji Trackmate plugin [10]. The obtained tracks were separated in three different fields: FRONT (< 50µm from the wound edge), MIDDLE (50µm < wound edge < 150µm) and BACK (150µm < wound edge < 300µm) according to the cell distance from the wound edge. Briefly, the distance between each nuclei centroid (identified by Trackmate analysis) and the wound edge (identified by the previously described wound healing analysis) was calculated using the Matlab “bwdist” function. The FRONT field corresponds to n ≤ 3 cells in a row.

For each section (FRONT, MIDDLE, BACK) and for each track the directional change rate (https://imagej.net/plugins/trackmate/algorithms#mean-directional-change) was measured by Fiji Trackmate plugin as the average angle difference between two subsequent displacements. The plotted directionality was then calculated as the inverse of the directional change rate.

**Kymograph analysis of cell protrusions at the wound edge**

For cell protrusion analysis at the wound edge, the wound healing assay was performed using Culture Inserts (Ibidi) to avoid debrides affecting the quality of the kymograph analysis. Inserts were placed in a 12-well plate and DCIS-RAB5A transfected cells were plated in each chamber (6x10^4^ cells/chamber). A cell-free wound area was created by removal of the insert. Cell migration was monitored by a Leica widefield Thunder imager equipped with a Leica sCMOS DFC9000GT, using a Leica HC PL Fluotar 20× objective, NA 0.5. Images were acquired every 30 seconds for 1 hour (100 ms exposure time). Images from 10 positions/condition were recorded every 30 seconds over a 1-hour period. To measure the dynamic of protrusive structures, each visible lamellipodia was analyzed by tracing a single pixel wide line orthogonally to the edge of the wound. The resulting kymograph was obtained and analyzed following the plugin of the ImageJ Kymograph macro (available at https://www.embl.de/eamnet/html/kymograph.html). Briefly, a segmented line was traced following the edges of the lamellipodia in the kymographs, and every protrusion or retraction in the video were averaged to obtain a single value for the plot. Persistence was calculated considering the total amount of frames in which the protrusion is visible until retraction begins.

In order to extract the lamellipodia region area from phase contrast time lapse images acquired as described above, we used an already described fully convolutional neural network [9]. The network was trained with 65 images of leading-edge cell protrusions at the wound.

**Cell sheet streaming and kinetic parameter measurements.**

DCIS-RAB5A and derivative cell lines were transfected and seeded as described for the wound healing experiment. RAB5A, DOCK7 and RAC1 WT/P29S expression was induced 16 hours before the experiment was initiated with 2.5 μg/mL doxycycline. The assay was performed using an environmental microscope incubator set to 37 °C and 5 % CO2 perfusion. An Olympus ScanR inverted microscope with 10× objective was used to acquire images every 5 minutes over a 24-hours period.

Quantification of monolayer dynamics in the streaming assays was performed using both Particle Imaging Velocimetry (PIV) and Particle Tracking (PT). PIV analysis of phase-contrast image sequences was performed using the Matlab software PIVlab [10]. In all cases, we adopted a final interrogation area of $20.8x20.8 \mu m^{2}$, close to the typical cell projected area. Spurious contributions to the velocity field corresponding to instantaneous global translations between consecutive frames (due to stage positioning errors) were removed through the smoothing procedure described in [11].

From the resulting velocity field $\boldsymbol{v}(\boldsymbol{x},t)$, where $\boldsymbol{x}$ is the position in the monolayer plane, the root mean squared velocity $v_{RMS}$ and the polar order parameter $\psi$ were computed as:

$$v_{RMS}=<\sqrt{\left\langle\left| \boldsymbol{v} \right|^{2} \right\rangle_{\boldsymbol{x}}}>_{j,t}$$

$$\psi=\left\langle\frac{\left| \left\langle\boldsymbol{v} \right\rangle_{\boldsymbol{x}} \right|^{2}}{\left\langle\left| \boldsymbol{v} \right|^{2} \right\rangle_{\boldsymbol{x}}} \right\rangle_{j,t}$$

Where $<{\ldots>}_{\boldsymbol{x}}$ denoted the space average while $<{\ldots>}_{j,t}$ denoted the average over different field of views (FOVs) $j$ and over different time points $t$. For each experiment, at least 5 different FOV are considered, while the time average was made over a time window from 4-20 hours after time-lapse recording starts. This time window roughly corresponded to the peak of RAB5A-induced motility[7].

The velocity spatial correlation function was evaluated from the PIV velocity field as

$$C_{vv}\left( r \right)=\left\langle\left\langle\frac{\left\langle\boldsymbol{v}\left( \boldsymbol{x},t \right)\cdot\boldsymbol{v}\left( \boldsymbol{x+r},t \right) \right\rangle_{\boldsymbol{x}}}{\left\langle\left| \boldsymbol{v}\left( \boldsymbol{x},t \right) \right|^{2} \right\rangle_{\boldsymbol{x}}} \right\rangle_{\theta} \right\rangle_{j,t}$$

Where $\left\langle\ldots\right\rangle_{\theta}$ denoted the azimuthal average over different directions of space vector $\boldsymbol{r}$. The obtained velocity space correlation was fitted with a stretched exponential function $C_{vv}\left( r \right)=\exp\left[ -\left( \frac{r}{l} \right)^{\beta} \right]$ from which the correlation length $L_{corr}=\frac{l}{\beta}\Gamma\left( \frac{1}{\beta} \right)$ (where $\Gamma$ was the gamma function) was recovered.

In order to quantify the relative motion between neighboring cells, we performed particle tracking on fluorescent nuclei. The particle tracking algorithm is described in detail in [11]. Briefly, we performed for each frame $I$ a seeded watershed transformation of the gradient image $\nabla I$ to segment each fluorescent nucleus $k$ and compute its centers of mass $\boldsymbol{x}_{k}$, assumed to coincide with the geometrical center of mass of its projection on the plane. Single nuclei trajectories were then obtained by linking nuclei positions in subsequent frames using the available Matlab implementation by D. Blair and E. Dufresne (http://physics.georgetown.edu/matlab) of the Grier Crocker tracking algorithm[12]. From nuclear trajectories, the instantaneous velocity $\boldsymbol{v}_{INS,k}$ was then evaluated as $\boldsymbol{v}_{INS,k}\left( t \right)=\left[ \boldsymbol{x}_{k}\left( t+\Delta t \right)-\boldsymbol{x}_{k}(t) \right]/\Delta t$, where $\Delta t$ is the time between two acquired frames. In order to reduce tracking noise, we computed a weighted moving average $\boldsymbol{v}_{k}\left( t \right)$ by convolving the instantaneous velocity with a Gaussian filter of width 10 min.

We calculated the root mean squared relative velocity ${\Delta v}_{RMS}$ of two nuclei at distance $r$ as

$${\Delta v}_{RMS}\left( r \right)=\left\langle\sqrt{\frac{\int_{r-\delta r}^{r+\delta r} \sum_{k,k'} \left| \boldsymbol{v}_{k}-\boldsymbol{v}_{k^{'}} \right|^{2}\delta\left( r^{'}-\left| \boldsymbol{x}_{k}-\boldsymbol{x}_{k^{'}} \right| \right)dr'}{\int_{r-\delta r}^{r+\delta r} \sum_{k,k'} \delta\left( r^{'}-\left| \boldsymbol{x}_{k}-\boldsymbol{x}_{k^{'}} \right| \right)dr'}} \right\rangle_{j,t},$$

where $k$ and $k'$ run over all nuclei within the same FOV and the amplitude of the integration interval $[r-\delta r,r+\delta r]$ corresponds to $1.3$ micrometer. To obtain an estimate of the relative motion of adjacent cells, we evaluated by linear interpolation ${\Delta v}_{RMS}$ for $r=14 \mu m$, corresponding to the average distance between the center of mass of a nucleus and the ones of its first neighbors.

All of the kinetic parameters were evaluated separately for each independent experiment. Each data point in Figure 1E-F, in the central panel of Figure G, and in the left panel of Figure 1G corresponds to independent experiments.

**Measurements of cryptic lamellipodia dynamics**

Assays were performed as previously described [7]. Briefly, DCIS-RAB5A cells stably expressing EGFP-LifeAct were mixed in a 1:10 ratio with unlabeled DCIS-RAB5A cells and transfected and seeded as described for the wound healing experiment. Cell migration was monitored by time-lapse phase-contrast and fluorescence microscopy with an Olympus ScanR inverted microscope using a 20× objective and images were acquired every 90 seconds over a 6-hour period. The quantification of cryptic lamellipodia protrusion velocity was performed using the ADAPT plug-in of Fiji. Cryptic lamellipodia directionality was measured as the angle φ delimited by the direction of the single lamellipodium and the direction vector of the collective pack locomotion. φ = 0° indicates that protrusion and collective migration have the same direction; φ = 180° indicates that protrusion and collective migration have opposite directions. The assay was repeated five times for each condition and at least 25 cells/condition were counted for each experiment.

**Immunofluorescence (IF)**

DCIS-RAB5A cells were grown on coverslips and fixed with 4 % paraformaldehyde (PFA) for 10 minutes. Coverslips were incubated in PBS with 2 % BSA for 1 hour for blocking, followed by incubation with primary antibodies for 1 hour at room temperature (RT) (overnight at 4 °C in case of confluent cells) and secondary antibodies for 1 hour at RT in PBS containing 1 % BSA. Incubation with DAPI (Sigma-Aldrich, cat. D9542) for 10 minutes was performed to stain the nuclei. Coverslips were mounted on glass slides using Mowiol Mounting Medium (Calbiochem) and images were acquired using Leica TCS SP8 laser confocal scanner mounted on a Leica DMI 6000B inverted microscope equipped with HCX PL APO 63×/1.4 NA oil immersion objective.

For Figure 2B, quantification of the mean fluorescence signal of myosin VI across the cells in Z-stacks images was performed manually using Fiji. The assay was repeated four times and 15 cells/experiment were analyzed.

Colocalization analysis was carried out adapting the JaCOP FIJI4,5 plugin [13] and using a custom pipeline able to process multiple folders and multicolor images. Manders’ coefficients [14] were calculated considering phalloidin or myosin VI signal as image A and DOCK7 signal as image B. For myosin VI-DOCK7 colocalization (Figure 5C), the XZ resliced images of the z-stacks images were split into two parts to distinguish the contribution of the basal and the apical regions of the cells.

**FRET based RAC1/CDC42 activation assay**

DCIS-RAB5A cells stably expressing RAC1-2G [4] or CDC42-2G [5] FRET biosensors were mixed in a 1:10 ratio with unlabeled DCIS-RAB5A cells and transfected and seeded at confluency on coverslips. Three days after seeding, coverslips were fixed with 4 % PFA for 10 minutes. Images were acquired with GE HealthCare Deltavision OMX system, equipped with 2 PCO Edge 5.5 sCMOS cameras, using a 60× 1.42 NA Oil immersion objective.

A custom Fiji plugin was developed to calculate single cell FRET signal. Briefly, YPF channel was used to identify single cell edge allowing background removal (https://imagej.net/plugins/rolling-ball-background-subtraction) and Gaussian filter. The single cells were segmented using the ImageJ Li’s threshold method (https://imagej.net/plugins/auto-threshold#li). We considered as periphery the region extending 1.5 µm from the cell edge. The ratio between the FRET channel and the CFP channel was calculated using a python script to obtain the distribution parameters.

**Protein** **expression and purification**

Recombinant proteins were expressed in E. coli BL21 (DE3) at 18 °C for 16 hours after induction with 0.5 mM IPTG at an OD600 of 0.6.

For GST-fusion proteins, cell pellets were resuspended in lysis buffer (50 mM Hepes pH 7.5, 200 mM NaCl, 1 mM EDTA, 0.1 % NP40, 5 % glycerol, 0.1 mM PMSF, and 1:500 protease inhibitor cocktail (Calbiochem). Sonicated lysates were cleared by centrifugation at 16,000 rpm for 45 minutes at 4 °C. Supernatants were incubated with 1 ml of glutathione-sepharose beads (Cytiva) per liter of bacterial culture for 4 hours at 4 °C. Beads were washed four times with lysis buffer followed by four washes with high salt buffer (20 mM Tris-HCl pH 8.0, 1 M NaCl, 1 mM EDTA, 1 mM DTT, and 5 % glycerol) and finally equilibrated in cleavage buffer (20 mM Tris-HCl pH 8.0, 200 mM NaCl, 1 mM DTT, and 5 % glycerol). PreScission protease was added at a 1:50 (w/w) ratio (protease:substrate) and incubated for 16 hours at 4 °C. Cleaved proteins were concentrated in Amicon Ultra Centrifugal Filters (MW cut-off 10 and 30 kDa, respectively) (Merck Millipore) and loaded onto a Superdex 75 10/300 GL column (Cytiva) equilibrated with 20 mM Tris-HCl pH 8.0, 150 mM NaCl, 5 % glycerol, and 1 mM DTT. Fractions containing purified proteins were collected and concentrated.

For HisMBP-fusion DHR2 (DOCK7), cell pellets were lysed in 50 mM Na-Phosphate buffer pH 7.5, 200 mM NaCl, 10 mM imidazole, and 5 % glycerol. After sonication and clearance of the lysate by centrifugation, supernatants were incubated with 1 ml Ni-NTA agarose beads (Qiagen) per 1 litre of bacterial culture. Beads were washed four times with lysis buffer followed by four washes with high salt/imidazole buffer (20 mM Na-Phosphate buffer pH 7.5, 1 M NaCl, 30 mM imidazole, and 5 % glycerol). Proteins were then eluted from beads with buffer containing 20 mM Tris-HCl pH 8.0, 200 mM NaCl, 300 mM imidazole, and 5 % glycerol, and dialyzed over night at 4 °C in the same buffer without imidazole. Dialyzed proteins were concentrated and purified by SEC as described for the GST-fusion proteins.

**Co-Immunoprecipitation and Pull-down assays**

For co-immunoprecipitation (co-IP) analysis, 1 mg of fresh lysates were incubated with specific antibodies for 2 hours at 4 °C. Cells were lysed in JS buffer (50 mM Hepes, pH 7.5, 150 mM NaCl, 1.5 mM MgCl2, 5 mM EGTA, 10% glycerol, and 1% Triton X-100) supplemented with 20 mM sodium pyrophosphate, pH 7.5, 250 mM sodium fluoride, 2 mM PMSF, 10 mM sodium orthovanadate, and protease inhibitors (Calbiochem), and lysates were cleared by centrifugation.

For pull-down experiments, 500 μg of transfected HEK293T cellular lysates were incubated with 1 μM of GST-fusion proteins immobilized onto GSH beads for 2 hours at 4 °C in JS buffer. After extensive washes with JS buffer, beads were re-suspended in Laemmli buffer and proteins were analyzed through sodium dodecyl sulphate–polyacrylamide gel electrophoresis (SDS-PAGE, 4–20% TGX precast gel, Bio-Rad). Detection was performed either by staining the gels with Coomassie or by immunoblotting using specific antibodies.

For the evaluation of direct binding, recombinant GST-fusion proteins (MYO6 tail, MyUb-CBD and RAC1, respectively) immobilized onto beads were incubated with purified HisMBP-tagged DHR2 domain of DOCK7 (15 µM final concentration) for 3 hours at 4 °C in low salt buffer (20 mM Tris-HCl pH 8.0, 50 mM NaCl, 5% glycerol, and 1 mM DTT). Beads were washed four times with the same buffer supplemented with 1% Triton-X and the samples were run on SDS-PAGE. Detection was performed by immunoblotting using anti-His antibody. Ponceau-stained membranes were used to show equal loading.

The GTP-CRIB assay was performed as described in [2]. Briefly, 500 μg of cell lysates from DCIS-RAB5A mock or myosin VI depleted (KD) cell monolayers were incubated with purified GST-CRIB (5 µM final concentration) for 1 hour at 4 °C. Beads were washed five times with lysis buffer and subjected to SDS-Page followed by immunoblotting with anti-RAC1 or CDC42 antibodies. Quantification was performed by normalizing the intensities of the bands to the total amount of RAC1 in the lysates. Data are reported as fold change with respect to RAC1-GTP levels in the corresponding mock sample for each experiment. Five independent experiments were performed.

**AlphaFold2-Multimer prediction**

The computational resources of the High-Performance Computing Biowulf cluster of the NIH (http://hpc.nih.gov) was used to run AlphaFold2-Multimer. The top-ranking structure from 50 predicted structures was selected for visualization and display. Structures were analyzed and figures generated by using PyMol (PyMOL Molecular Graphics System, http://www.pymol.org).

**GEF activity assay**

Before starting the assay, all recombinant purified proteins were dialyzed in buffer containing 20 mM Tris-HCl pH 7.0, 150 mM NaCl and 10 mM MgCl2.

Recombinant RAC1 (10 µM final concentration) was pre-incubated with GDP (15 µM final concentration) for 30 minutes on ice in buffer containing 20 mM Tris-HCl pH 7.0, 150 mM NaCl, 10 mM MgCl2, and 0.2 mg/ml BSA. Subsequently, fluorescent boron-dipyrromethene-fluor (BODIPY-FL)-GTP (2.4 mM final concentration) was added to the GDP-loaded RAC1. To detect intrinsic RAC1 activity, buffer was added to the reaction prior to measurement. As positive control we used EDTA (12 mM final concentration). In the testing conditions, recombinant HisMBP-tagged DHR2 DOCK7 domain was added at the indicated final concentration. To evaluate myosin VI activity, recombinant MYO6 tail construct at the indicated concentration was pre-incubated on ice for 30 minutes with HisMBP-tagged DHR2 DOCK7 and then added to the reaction mixture. Kinetic hydrolysis of BODIPY-FL-GTP was measured at 30 °C by monitoring the increase in fluorescence at excitation/emission wavelengths of 485/535 nm in a black 384-well microplate (Corning) using EnVision (PerkinElmer) plate reader. A reaction containing buffer, GDP and BODIPY-FL-GTP at the same final concentration was set up in order to subtract background value. All reactions were performed in technical triplicates. At the end of the measurement, samples were run on SDS-PAGE and analyzed by Coomassie staining. The assay was repeated at least three times/condition.

**IHC analysis and quantification**

Formalin-fixed and paraffin embedded (FFPE) tissue samples of human breast cancer cases (8 ductal *in situ* carcinoma and 8 invasive ductal carcinoma cases, collected and handled according to the Helsinki Declaration) were selected for the quantitative *in situ* immunophenotypical analyses. Four-micrometers-thick tissue sections were deparaffinized, rehydrated and unmasked using Novocastra Epitope Retrieval Solutions pH6 in thermostatic bath at 98 °C for 30 minutes. Subsequently, the sections were brought to room temperature and washed in PBS. After neutralization of the endogenous peroxidase with 3 % H_2_O_2_ and Fc blocking by 0.4 % casein in PBS (Novocastra, Leica Microsystems), the samples were incubated for 90 minutes at room temperature with anti-myosin VI primary antibody (code #1296, homemade, 1:250). IHC staining was revealed using Novolink Polymer Detection Systems (Novocastra, Leica Microsystems) and Romulin AEC Chromogen kit (BioOptica) as substrate chromogen. Slides were counterstained with Harris hematoxylin (Novocastra, Leica Microsystems) and analyzed under a Zeiss Axioscope A1 microscope. Microphotographs were collected using a Zeiss Axiocam 503 Color digital camera with the Zen 2.0 Software (Zeiss). Quantitative analyses of IHC staining were performed by calculating the average percentage of positive signals in five nonoverlapping fields at medium-power magnification (200X) using the Positive Pixel Count v9 (2+ moderate positivity and 3+ strong positivity) ImageScope software. Average percentage of positive signals (3+ or 2+) was calculated in five nonoverlapping fields of view (200x). This study was approved by the University of Palermo Ethical Review Board (approval number 09/2018).

**Expression profile analysis**

The transcriptional myosin VI isoforms expression was analyzed by using TSVdb (http://www.tsvdb.com)[15], using RNA-seq data level of The Cancer Genome Atlas (TCGA) breast cancer (BRCA) dataset which includes 981 invasive breast carcinoma samples (and 112 normal breast samples) with complete clinical and pathological information together with PAM50 and SigClust Subtype assignments. Briefly, we downloaded the normalized expression level (RPKM) of short (isoform_uc003pii) and long (isoform_uc003pih) myosin VI isoforms mapped by TSVdb on GRCh37/hg19 genome assembly. RPKM data were +1 trimmed to avoid “0” or “Null” values before processing these to obtain ratios of expression between short versus long myosin VI isoforms. Statistical analyses and relative plots were done using JMP 17 (SAS) software.

**Key resources table**

| **REAGENT or RESOURCE** | **SOURCE** | **IDENTIFIER** |
| --- | --- | --- |
| Antibodies | | |
| anti-Desmoplakin | provided by Kathleen Green and Lisa Godsel | N/A |
| anti-RAC1 | Becton Dickinson | Cat#610651 |
| anti-MYO6 | Homemade[6] | Eurogentec-1296 |
| anti-GAPDH | Santa Cruz | Cat#32233 |
| anti-DOCK7 | Santa Cruz | Cat#398888 |
| anti-RAB5A | Santa Cruz | Cat#166600 |
| anti-CDC42 | Santa Cruz | Cat#390210 |
| anti-GFP | Sigma-Aldrich | Cat#G1544 |
| anti-His | Cell Signaling | Cat#2366 |
| Chemicals, peptides, and recombinant proteins | | |
| Human EGF | Vinci Biochem | BPS-90201-1 |
| GST-beads | GE Healthcare | GE17-0756-01 |
| GFP-TRAP | Cromotek | RRID: AB_2631357 |
| BODIPY™ FL-GTP | Thermo Fisher Scientific | Cat# G12411 |
| Carbachol | Calbiochem, Sigma | CAS 51-83-2 |
| Experimental models: Cell lines | | |
| MCF10.DCIS.com RAB5A | [7] | N/A |
| MCF10.DCIS.com RAB5A GFP-LifeAct mCherry-H2B | [7] | N/A |
| MCF10.DCIS.com RAB5A mCherry-H2B | [7] | N/A |
| MCF10.DCIS.com RAB5A GFP-DOCK7 | This study | N/A |
| MCF10.DCIS.com RAB5A GFP-LifeAct mCherry-H2B RAC1 WT | This study | N/A |
| MCF10.DCIS.com RAB5A GFP-LifeAct mCherry-H2B RAC1 P62S | This study | N/A |
| MCF10.DCIS.com RAB5A RAC1-2G | This study | N/A |
| MCF10.DCIS.com RAB5A CDC42-2G | This study | N/A |
| HEK293T | ICLC | N/A |
| Oligonucleotides | | |
| siRNA targeting sequence: MYO6 #1: 5′-GAGGCUGCACUAGAUACUUUGCUAA-3′ | Thermo Fisher Scientific | NM_004999_stealth_1106 |
| siRNA targeting sequence: MYO6 #2: 5′-GAGCCTTTGCCATGGTACTTAGGTA-3′ | Thermo Fisher Scientific | NM_004999_stealth_4 |
| siRNA targeting sequence: DOCK7 #1: 5′- UUUAAGGUCAUCUUGAUCAUCCUGG-3′ | Thermo Fisher Scientific | Cat # HSS131697 |
| siRNA targeting sequence: DOCK7 #2: 5′- AUUAGGGUAAGUAUUUGGUAGGCGG-3′ | Thermo Fisher Scientific | Cat # HSS131695 |
| MYO6 isoform detection by PCR For. Sequence: CCGAGCTCATCAGTGATGAGGC | [6] | N/A |
| MYO6 isoform detection by PCR Rev. Sequence: CCAAGCATGATACACTTTTAGTCTCC | [6] | N/A |
| Primer: DOCK7-FL. Forward:  GCGGCCCCGAACTAGTGCCACCATGGTGAGCAAGG | This Paper | N/A |
| Primer: DHR2-LobeBC (aa1795-2072) of DOCK7. EcoRI-Forward: AAAGAATTCCGGATGTTTGGCACCTATTTTC | This Paper | N/A |
| Primer: DHR2-LobeBC (aa1795-2072) of DOCK7. XhoI-Reverse: AAACTCGAGTTAAGGGATCTTTCTGTTGATC | This Paper | N/A |
| Primer: DHR2-LobeAC (aa1597-2072) of DOCK7. EcoRI-Forward: AAAGAATTCGATCTGGTTTTCAATCTCC | This Paper | N/A |
| Primer: DHR2-LobeAC (aa1597-2072) of DOCK7. XhoI-Reverse: AAAGAATTCAAGGGTTACCAGACCTCTCC | This Paper | N/A |
| Primer: RAC1 (aa1-177). EcoRI-Forward: AAAGAATTCATGCAGGCCATCAAGTG | This Paper | N/A |
| Primer: RAC1 (aa1-177). XhoI-Reverse: AAACTCGAGTTAGAGGACTGCTCG | This Paper | N/A |
| Recombinant DNA | | |
| pLenti-RAC1-2G FRET Biosensor | [4] | Addgene Plasmid #66111 |
| pLenti-Cdc42-2G FRET Biosensor | [5] | Addgene Plasmid #68813 |
| pGEX-GST-CRIB | [16] | N/A |

**SUPPLEMENTARY FIGURES**

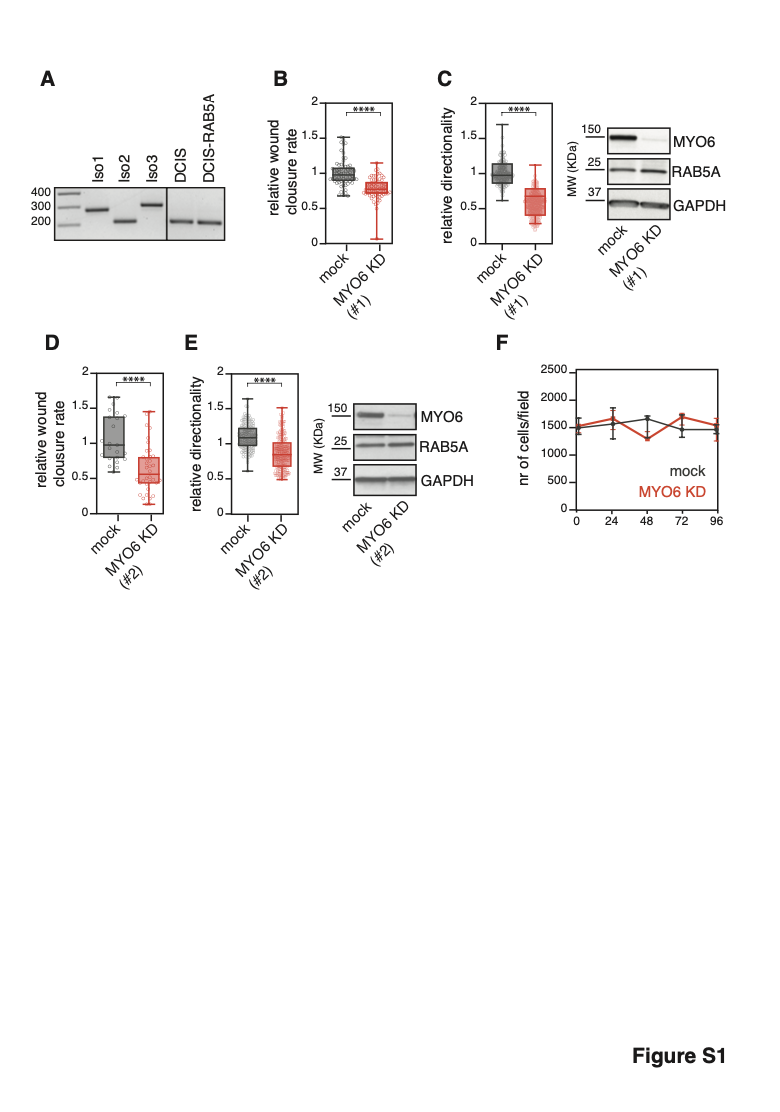

**Figure S1. Myosin VI is critical for wound closure.**

**(A)** RT-PCR analysis of myosin VI isoforms of DCIS and DCIS-RAB5A cells grown at confluency for three days. Oligos used for the PCR map in Exon 28 and Exon 33 and controls are from plasmids carrying the tails of the different isoforms as in [31]. Myosin VI short (isoform 2) is the only isoform expressed in DCIS cell lines.

**(B-E**) Wound healing assay. DCIS-RAB5A cells expressing H2B-mCherry were transfected in suspension with the indicated siRNA oligos, seeded at confluency in six-well plates (1.5x10^6^ cells/well), transfected again the day after and then cultured for an additional day to obtain a uniform monolayer. 16 hours before the scratch, 2.5 μg/mL doxycycline was added to fresh media to induce the expression of RAB5A. After the scratch, the closure of the wound was monitored by time-lapse microscopy. See also Video S3.

**(B)** Motility rate was quantified by measuring the wound-closure area covered over time. The average wound closure speed relative to mock condition is plotted in the graph. Empty circle represents the mean wound closure velocity quantified for each video. *n*≅60 (at least 10 videos/condition for six independent experiments. ****P < 0.0001 by Student’s t-test.

**(C)** Cell directionality was quantified by tracking the motion of mCherry-positive nuclei with Fiji TrackMate plugin. The plotted directionality is the inverse of the directional change rate parameter obtained with TrackMate analysis. Results in the graphs are expressed relative to mock average value. Empty circle represents the mean directionality of cell tracks. *n*≅180 (at least 45 cells/condition for four independent experiments. ****P < 0.0001 by Student’s t-test. Right panel, IB analysis of the myosin VI depleted lysates performed at the end of the experiment.

**(D**, **E**) Same experiments described in B and C, but using siRNA#2 oligo. In D, Empty circle represents the mean wound closure velocity quantified for each video. *n*≅30 (at least 5 videos/condition for four independent experiments. In E, Empty circle represents the mean directionality of cell track. *n*≅90 (at least 30 cells/condition for three independent experiments. ****P < 0.0001 by Student’s t-test. Right panel, IB analysis of the myosin VI depleted lysates performed at the end of the experiment.

**(F)** Proliferation curves of mock or myosin VI depleted (KD) DCIS-RAB5A cell monolayers monitored by time-lapse phase contrast microscopy for fours days. Every 24 hours cells were counted and expressed as number of cells/field of view. Data are the mean ± SD (*n*=3 from at least five fields/condition for three independent experiment).

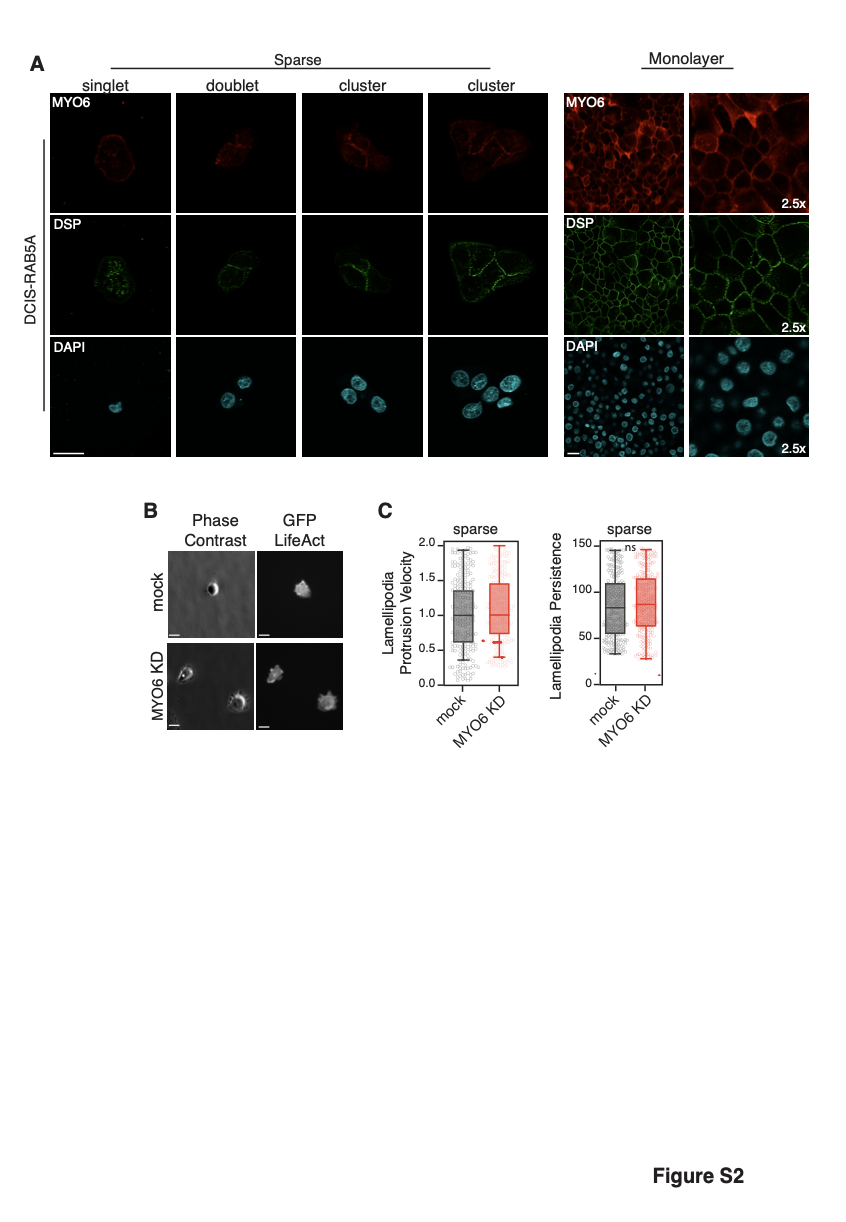

**Figure S2. Characterization of myosin VI localization and activity.**

**(A)** IF analysis of singlet, doublet, cell clusters (left panels) or monolayer (right panel) of DCIS-RAB5A cells. Middle plane is shown. Red, myosin VI; green, Desmoplakin staining (DSP); cyan, DAPI. Scale bar, 25μm. Myosin VI shows a diffuse cytoplasmic pattern at single-cell level and a rapid accumulation at desmosomes when a cluster of two or more cells is established.

**(B)** Phase-contrast and fluorescent images of lamellipodia of DCIS-RAB5A cells expressing GFP-LifeAct, mock or myosin VI depleted (KD), seeded in sparse conditions.

**(C)** Quantification of lamellipodia protrusion velocity (left panel) and persistence (right panel). Images of lamellipodia from time-lapse microscopy were quantified using ADAPT plug-in of Fiji and velocity results are expressed relative to GFP-LifeAct-expressing mock average value. Persistence is plotted as the number of frames in which the lamellipodia is detectable, from its appearance to its disappearance. Left panel, empty circle represents the mean lamellipodia protrusion velocity of a single cell. *n*≅200 (at least 50 cells/condition for four independent experiments). Right panel, empty circle represents the persistence of single lamellipodia. *n*≅250 (at least 60 lamellipodia/condition for four independent experiments). ns > 0.999 by Student’s t-test.

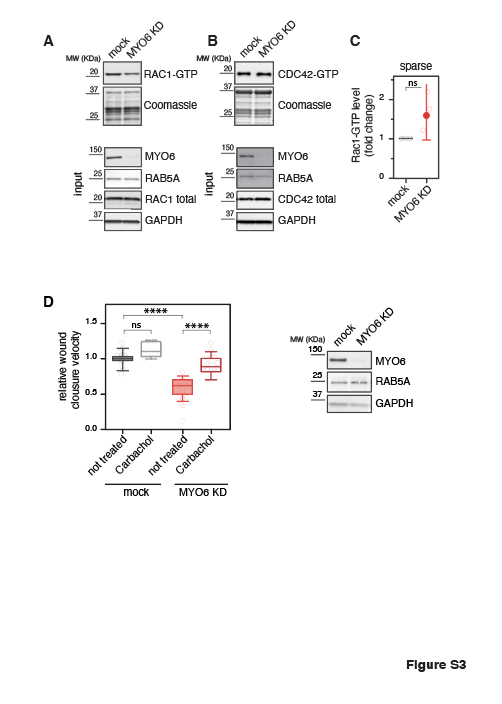

**Figure S3. Myosin VI depletion reduces RAC1 GTPase activation in cell monolayers.**

**(A)** Example of GST-CRIB assay performed to evaluate RAC1-GTP level in lysates from DCIS-RAB5A mock or myosin VI depleted (KD) monolayers. IB as indicated. Ponceau shows equal loading of the GST proteins.

**(B)** Example of GST-CRIB assay performed to evaluate CDC42-GTP level in lysates from DCIS-RAB5A mock and myosin VI depleted (KD) cell monolayers. IB as indicated. Ponceau shows equal loading of the GST proteins.

**(C)** Quantification of the GST-CRIB assay performed using lysates from DCIS-RAB5A mock or myosin VI depleted (KD) cells seeded in sparse condition. The intensity of the RAC1-GTP band was normalized to the total amount of RAC1 present in the lysate. Data are reported as fold change with respect to RAC1-GTP level in the corresponding mock sample for each experiment. Values are means ± SD. *n*=5 independent experiments. Results in the graph are expressed relative to mock average value. ns > 0.999 by Student’s t-test.

**(D)** Wound healing assay of DCIS-RAB5A mock and myosin VI depleted (KD) cells treated with Carbachol (Calbiochem, Sigma) at 100 nM concentration added after wound generation. Motility was quantified by measuring the area covered over time. The average wound closure speed relative to mock condition is plotted in the graph. Right panel, IB analysis with the indicated antibodies of the lysates obtained at the end of the experiment. Empty circle represents the mean wound closure velocity quantified for each video. *n*≅12 (at least four videos/condition for three independent experiments. ns > 0.999, **P < 0.01, ****P < 0.0001 by ANOVA test.

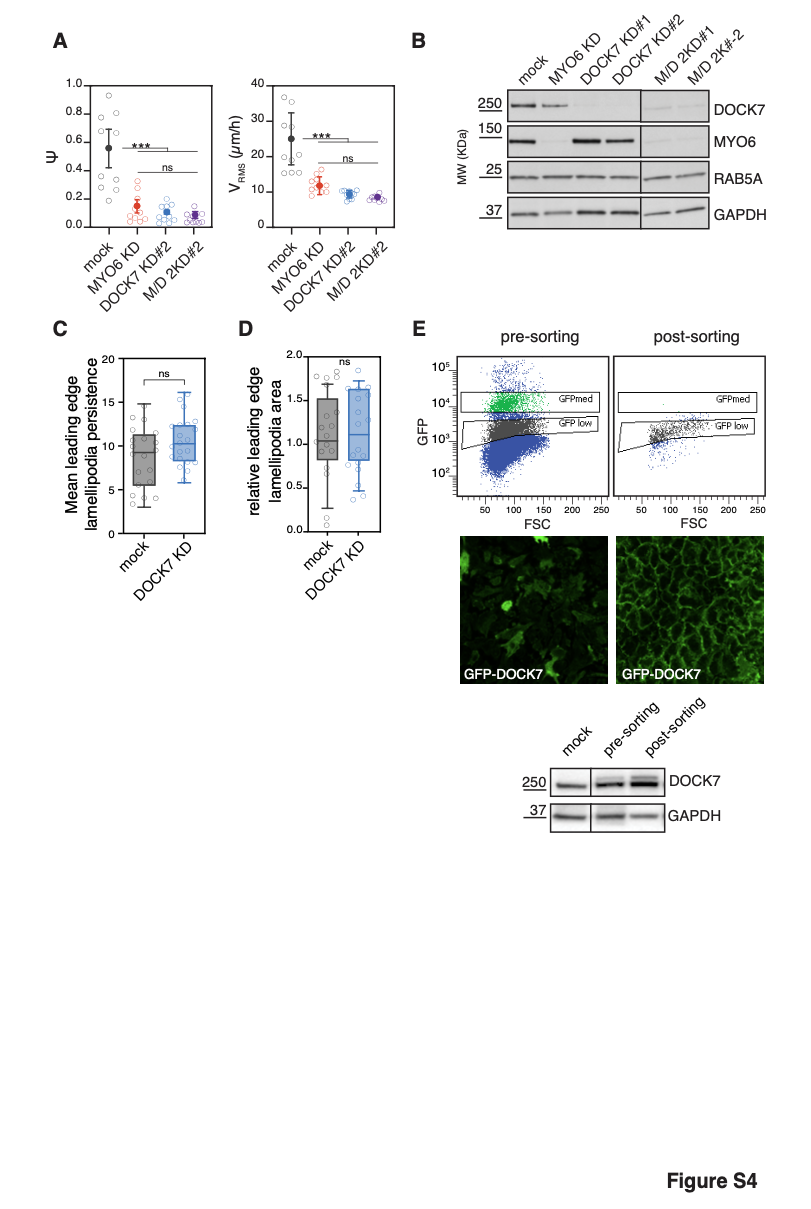

**Figure S4. Role of DOCK7 activity at cryptic lamellipodia.**

**(A)** PIV analysis of the streaming of DCIS-RAB5A mock, MYO6 and DOCK7 depleted singularly or in combination (M/D 2KD) cells as in Figure 4D using a second siRNA oligo for DOCK7 (#2). From left to right: orientational order parameter $\psi$ and root mean squared velocity $v_{RMS}$. Empty circle represents mean of each experiment calculated from at least three videos/condition for three independent experiments. Error bars ± SD. ns > 0.999, ***P < 0.001 by ANOVA test.

**(B)** IB analysis of DCIS-RAB5A cells lysed at the end of the experiments shown in Figure 4D and Figure S4A with the indicated antibodies. Both siRNA oligos against DOCK7 are efficient in depleting the protein both when used alone or in combination with myosin VI siRNA oligos.

**(C)** Quantification of leading edge lamellipodia persistence (expressed in minutes) obtained by kymograph analysis of GFP-LifeAct expressing DCIS-RAB5A mock or DOCK7 depleted (KD) scratched cell monolayers. described in C. Empty circle represents the persistence of single leading edge lamellipodia. *n*≅20 (at least 5 lamellipodia/condition for three independent experiments). ns > 0.999 by Student’s t-test.

**(D)** Quantification of leading edge lamellipodia mean area by a fully convolutional neural network. Phase-contrast images of wounded DCIS-RAB5A mock or DOCK7 depleted (KD) cell monolayers were used. Results in the graph are expressed relative to mock average value. Empty circle represents the area occupied by leading edge lamellipodia. *n*≅20 (at least 5 lamellipodia/condition for four independent experiments). ns > 0.999 by Student’s t-test.

**(E)** FACS analysis of DCIS-RAB5A expressing pSLIK-GFP-DOCK7, before and after the sorting procedure. Bottom panel, from left to right: representative IF images and IB analysis of the same populations with the indicated antibodies.

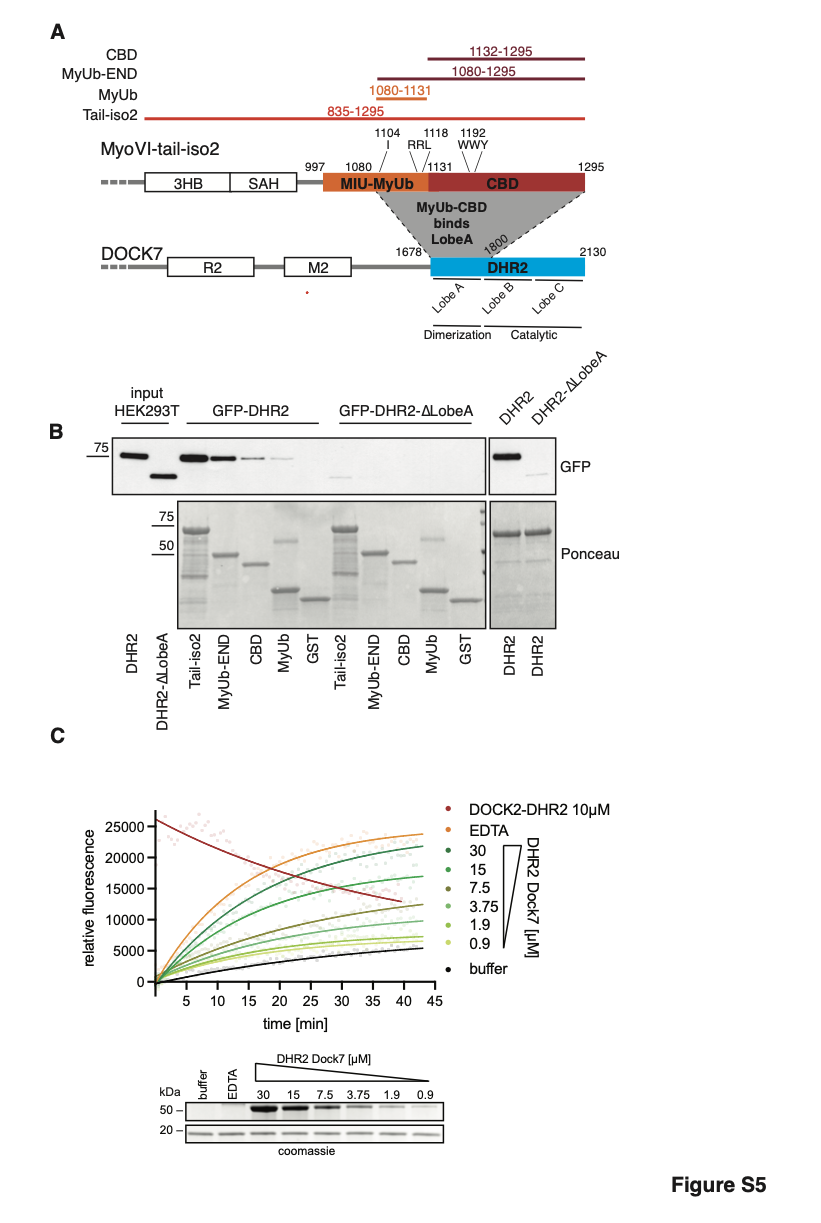

**Figure S5. Mapping of the interaction surface between DOCK7 and MYO6.**

**(A)** Scheme of DOCK7 and MYO6 tail domain organization, different GST-tagged myosin VI constructs used in the pull-down experiment and boundaries of the lobe A,B and C of the DHR2 of DOCK7.

**(B)** GST pull-down assay using the indicated myosin VI constructs and lysates from HEK293T cells transfected with GFP-DHR2 or GFP-DHR2-ΔLobeA of DOCK7. Bound proteins were resolved in SDS-PAGE and analyzed by IB with anti-GFP antibody. Ponceau as indicated.

(**C**) Representative RAC1 GEF activity assay using the indicated concentration of the DHR2 domain of DOCK7 and DOCK2 for comparison. Bottom, Coomassie gel as loading control.

**Video S1. Myosin VI depletion does not affect random migration of single cell.**

Time-lapse images of single cell motility of the indicated cell lines. Top, red (H2B-mCherry) channel. Bottom, phase-contrast images with the track of the followed nuclei over time. Cells were monitored over a 20-hour period, with a frame rate of 5 minutes. Scale bar, 100 µm.

**Video S2. Myosin VI depletion does not affect confined migration of single cell.**

Time-lapse images of single cell motility of the indicated cell lines seeded on a line micro-pattern (10 μm diameter). Left, phase-contrast images. Right, red (H2B-mCherry) channel. Cells were monitored over a 24-hour period, with a frame rate of 5 minutes. Scale bar, 100 µm.

**Video S3. Myosin VI depletion affects collective cell migration.**

Time-lapse images of wound healing assay of the indicated cell lines. Cells were monitored over a 24-hour period, with a frame rate of 5 minutes. The videos were stopped at wound closure. Scale bar, 100 µm.

**Video S4. Myosin VI depletion affects collective cell streaming.**

Time-lapse images of streaming assay of the indicated cell lines. Cells were monitored over a 24-hour period, with a frame rate of 5 minutes. Scale bar, 100 µm.

**Video S5. Myosin VI depletion affects directional collective cell migration.**

Video of the velocity field (vector map) obtained by PIV analysis of streaming assay. Cells were monitored over a 24-hour period. The color map represents the alignment with respect to the mean instantaneous velocity, quantified by the alignment index a(x)=[v(x)⋅v_0]/(|v(x)||v_0 |) that is equal to 1(−1) when the local velocity is parallel (antiparallel) to the mean direction of migration. The corresponding mean velocity is shown in the inset of each panel.

**Video S6. Myosin VI depletion does not affect the formation of cryptic lamellipodia.**

Time-lapse images of fluorescent (Top) and phase-contrast (Bottom) DCIS-RAB5A cell monolayers where GFP-LifeAct-expressing mock or myosin VI depleted cells were interspersed (1:10 ratio). Cells were monitored over an 8-hour period, with a frame rate of 90 seconds. Scale bars, 15 μm.

**Video S7. RAC1 WT and P29S mutant overexpression rescue wound healing defects caused by myosin VI depletion.**

Time-lapse images of wound healing assay of the indicated cell lines. Cells were monitored over a 24-hour period, with a frame rate of 5 minutes. The videos were stopped at wound closure. Scale bar, 100 µm.

**Video S8. RAC1 WT overexpression rescues streaming defects caused by myosin VI depletion.**

Time-lapse images of streaming assay of the indicated cell lines. Cells were monitored over a 24-hour period, with a frame rate of 5 minutes. Scale bar, 100 µm.

**Video S9. DOCK7 depletion affects collective cell migration.**

Time lapse images of wound healing assay of the indicated cell lines. Cells were monitored over a 24h period, with a frame rate of 5 minutes. The videos were stopped at wound closure. Scale bar, 100 µm.

**Video S10. DOCK7 depletion affects collective cell streaming.**

Time lapse images of streaming assay of the indicated cell lines. Cells were monitored over a 24-hour period, with a frame rate of 5 minutes. Scale bar, 100 µm.
